# The multivalent G-quadruplex (G4)-ligands MultiTASQs allow for versatile click chemistry-based investigations

**DOI:** 10.1101/2022.10.28.512542

**Authors:** Francesco Rota Sperti, Jérémie Mitteaux, Joanna Zell, Angélique Pipier, Ibai E. Valverde, David Monchaud

## Abstract

Chemical biology hinges on multivalent molecular tools that uniquely allow for interrogating and/or manipulating cellular circuitries from the inside. The success of many of these approaches relies on molecular tools that make it possible to visualize biological targets in cells and then isolate them for identification purposes. To this end, click chemistry has become in just a few years a vital tool in offering practically convenient solutions to address highly complicated biological question. We report here on two clickable molecular tools, the biomimetic G-quadruplex (G4) ligands MultiTASQ and ^az^MultiTASQ, which benefit from the versatility of two types of bioorthogonal chemistry, CuAAC and SPAAC (the discovery of which was very recently awarded the Nobel Prize of chemistry). These two MultiTASQs are here used to both visualize G4s in, and identify G4s from human cells. To this end, we developed click chemo-precipitation of G-quadruplexes (G4-click-CP) and *in situ* G4 click imaging protocols, which provide unique insights into G4 biology in a straightforward and reliable manner.

## I. Introduction

Click chemistry, either the copper(I)-catalyzed azide-alkyne cycloaddition (CuAAC) developed by Meldal^1^ and Sharpless^2^ or its metal-free counterpart strain-promoted azide-alkyne cycloaddition (SPAAC) developed by Bertozzi,^3^ finds wide applications in chemistry and chemical biology, as recognized very recently by the Nobel committee.^4^ Many bioorthogonal strategies aiming at interrogating cell circuitries with molecular modulators now hinge on click chemistry: for example, click chemistry is widely used for imaging purposes^5^ in both fixed (CuAAC) and live cells (SPAAC); it is also used for pulling down probes in interaction with their cellular partners and/or the genomic targets followed by either proteomics (‘click pull-down’ or ‘click-proteomics’) or sequencing (‘click-seq’ or ‘chem-click-seq’). An illustrative example is the clickable analog of Remodelin,^6^ which was clicked *in situ* to AF488-azide for localization purposes in human osteosarcoma (U2OS) cells, and to a biotin-azide derivative to identify the acetyl-transferase NAT10 as its cellular partners.^6^ Similarly, a clickable analog of the BET inhibitor JQ1^7^ termed JQ1-PA was labeled *in situ* with AF488-azide for localization purposes in human leukemia (MV4;11) cells, and to a biotin-azide derivative to identify the genomic binding sites of bromodomain-containing protein 4 (BRD4), which is targeted by JQ1.^8^ Also, a series of clickable Olaparib^9^ derivatives were exploited to confirm the specificity of this drug to poly(ADP-ribose)-polymerase 1 (PARP1) in human cervical cancer (HeLa) cells, *via* a combination of click-imaging and click-proteomics.^10^

In the field of G-quadruplexes (G4s), the CuAAC allowed first and foremost for the modular synthesis of a wide variety of G4-ligands.^11^ When applied to bioorthogonal investigations, click chemistry has permitted the very first visualization of G4s in human cells, using a clickable pyridostatin (PDS)^12^ derivative termed PDS-α labeled *in situ* with AF594-azide in U2OS (by CuAAC),^13^ then in human colon cancer (HT-29) cells using clickable PhenDC3^14^ derivatives (PhenDC3-alk for CuAAC, PhenDC3-az for SPAAC)^15^ and again in U2OS with a clickable L2H2-6OTD^16^ derivative termed L2H2-6OTD-az (by SPAAC).^17^ Another approach referred to as G4-GIS (for G4-ligand guided immunofluorescence staining) involved a series of clickable pyridodicarboxamide (PDC) derivatives, notably PDC-4,3-Alk that was used either pre-clicked or *in situ* clicked with 5-BrdU-N_3_ (a 5-bromo-2’-deoxyuridine functionalized with an azide group) in human lung cancer (A549) cells, prior to be immunodetected using an anti-5-BrdU antibody. Two proteomics-based approaches termed G4-LIMCAP (for G4 Ligand-mediated cross-linking and pull-down)^18^ and co-binding-mediated protein profiling (CMPP),^19^ based on two other clickable PDS derivatives (PDB-DA-A, and photoPDS, respectively), were recently used to uncover several new G4-binding proteins in human breast cancer (MDA-MB-231) cells, immortalized human fibroblast (SV589) cells,^18^ and human embryonic kidney (HEK293T) cells.^19^

These examples brightly illustrate the interest of clickable probes in chemical biology in general, and in the G4 field in particular. Following up on our recent use of biotinylated G4-specific molecular probes BioTASQ, BioCyTASQ and BioTriazoTASQ (Figure 1A)^20–24^ to isolate G4s *via* affinity precipitation, we aim here at further exploiting the exquisite G4 selectivity of TASQs: this specificity originates in the biomimetic, like-likes-like interaction between the G-quartet of the G4 and the synthetic G-quartet of the TASQ (for template-assembled synthetic G-quartet, Figure 1B).^25^ TASQs are smart ligands that adopt their G4-affinic conformation only in presence of their G4 targets, which thus makes them uniquely actively selective for G4s. We thus report here on our new, patented MultiTASQ technology (Figure 1A),^26^ which comprises multivalent TASQs with either an alkyne appendage (MultiTASQ) for CuAAC applications or an azide chain (azidoMultiTASQ, or ^az^MultiTASQ) for SPAAC applications, used for both click chemo-precipitation of G-quadruplexes (G4-click-CP) and click-imaging purposes (Figure 1B).

**Figure 1.**
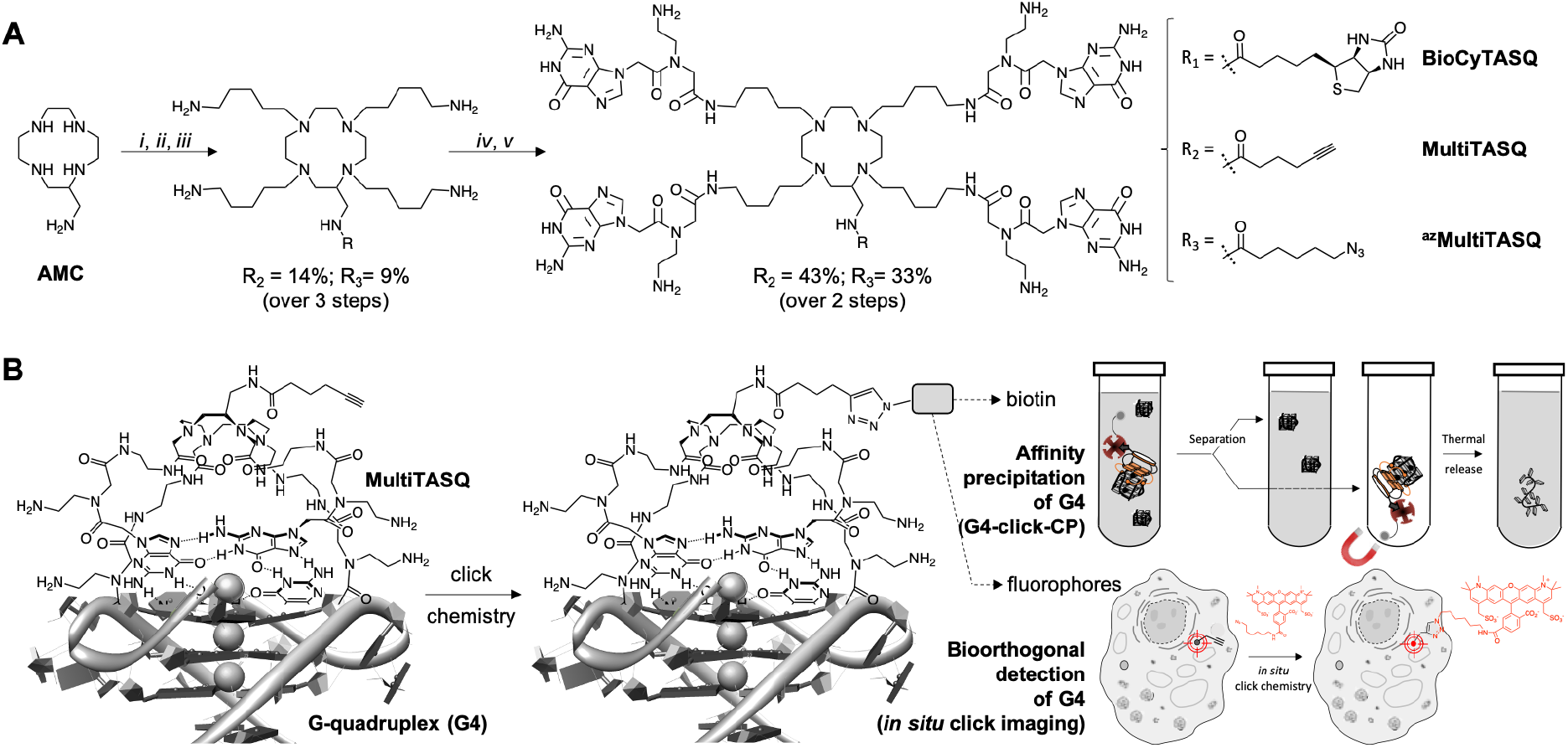
**A**. Chemical structure and synthesis of BioCyTASQ, MultiTASQ and ^az^MultiTASQ; *i*: R-CO_2_H, TSTU, DIPEA, DMF; *ii*: MsO-(CH_2_)_5_-NHBoc, K_2_CO_3_, CH_3_CN; *iii*: TFA; *iv*: Boc-^PNA^G-OH, HBTU, DIPEA, DMF; *v*: TFA. **B**. Schematic representation of the G-quartet/G-quartet interaction between the TASQ and a DNA/RNA G4, and of the click chemistry-based investigations made possible with MultiTASQs, *i.e*., G4 isolation by affinity capture (clicked biotin) and optical imaging (clicked fluorophore).

## II. Design and synthesis of MultiTASQs

The design of MultiTASQs was inspired by the recently developed biotinylated TASQs,^20–23^ with the goal of adding a greater degree of bio-compatibility and versatility. By changing the biotin appendage for an alkyne or an azide one, the resulting TASQ could be usable for live-cell incubation as the triple bond/azide minimally divert TASQ biodistribution^5^ (unlike biotin that can creates H-bonds with various cellular components), which makes this technology implementable in living cells.^27^ Furthermore, the inability of the alkyne/azide appendage to form H-bond will also preclude internal poisoning of the TASQ, found to be responsible of the lower G4-affinity of BioTASQ as compared to non-biotinylated TASQ.^21–22^

The synthetic pathway of MultiTASQs (Figures 1 and S1-S10) thus started from the aminomethylcylen (AMC)^28^ coupled with 5-hexynoic acid (MultiTASQ) or 6-azido-hexanoic acid (^az^MultiTASQ) to obtain compound the corresponding AMC derivatives in 38 and 21% chemical yield, respectively. These derivatives were subsequently reacted with an excess of 5-(Boc-amino)pentylmesylate linker (8.0 mol. equiv., 38 and 40% chemical yield, respectively), deprotected by TFA (quantitative) and engaged in reaction with Boc-^PNA^G-OH^29^ (4.4 mol. equiv.) to provide the Boc-protected MultiTASQs with 43 and 33% chemical yield, respectively. MultiTASQs were then deprotected prior to use with TFA, which led to the final compounds in a 6.2 and 2.7% chemical yield over 5 steps, respectively.

## III. *in vitro* validation of MultiTASQs

### G4 affinity and selectivity of MultiTASQs

The G4 affinity of the two MultiTASQs was evaluated *via* competitive FRET-melting assay^30^ against two DNA G4s (Table S1), from sequences found in human telomere (F21T) and in promoter region of the Myc gene (F-Myc-T), and two RNA G4s (Table S1), from sequences found in human telomeric transcript (F-TERRA-T) and in UTR region of the VEGF mRNA (F-VEGF-T). These experiments were performed with labelled DNA/RNA (0.2 μM) and TASQs (1 μM, 5 mol. equiv.) in the absence or presence of an excess of competitive dsDNA (calf thymus DNA, or CT-DNA, 15 or 50 mol. equiv.). Results seen in Figure 2A indicated that *i*- TASQs display a lower affinity for DNA G4s (ΔT_1/2_ = 2.5 ± 0.2 and 5.3 ± 0.5 °C for MultiTASQ, and 2.2 ± 0.3 and 6.4 ± 0.2 °C for ^az^MultiTASQ at 5 mol. equiv. ligand) as compared to RNA G4s (ΔT_1/2_ = 11.0 ± 0.9 and 11.9 ± 0.4 °C for MultiTASQ, and 12.4 ± 0.6 and 11.3 ± 0.4 °C for ^az^MultiTASQ at 5 mol. equiv. ligand), and *ii*- TASQs are extremely selective for G4s over dsDNA (averaged ^FRET^S > 1.0 for both TASQs at 50 mol. equiv. CT-DNA). These results were in line with what was obtained with the previously reported TASQs.^22–23^ Their G4-affinity was confirmed *via* an equilibrium-binding assay that relies on the use of a G4 (here, Myc, Table S2) labeled with a Cy5 dye (on its 5’-end) that is quenched upon ligand binding.^31^ This assay was calibrated with PhenDC3,^14^ as it was used in the initial setup (^app^K_D_ = 29.6 ± 0.9 nM); in our hands (Figure 2B), the G4 affinity of PhenDC3 was confirmed (^app^K_D_ = 57.7 ± 0.2 nM), and that of MultiTASQs lower (^app^K_D_ = 1.44 ± 0.1 and 0.90 ± 0.1 μM for MultiTASQ and ^az^MultiTASQ, respectively) and in line with that of BioCyTASQ (^app^K_D_ = 1.01 ± 0.1 μM). These results thus showed that the modification of the TASQ’s appendage did not modify their affinity and selectivity for G4s.

**Figure 2.**
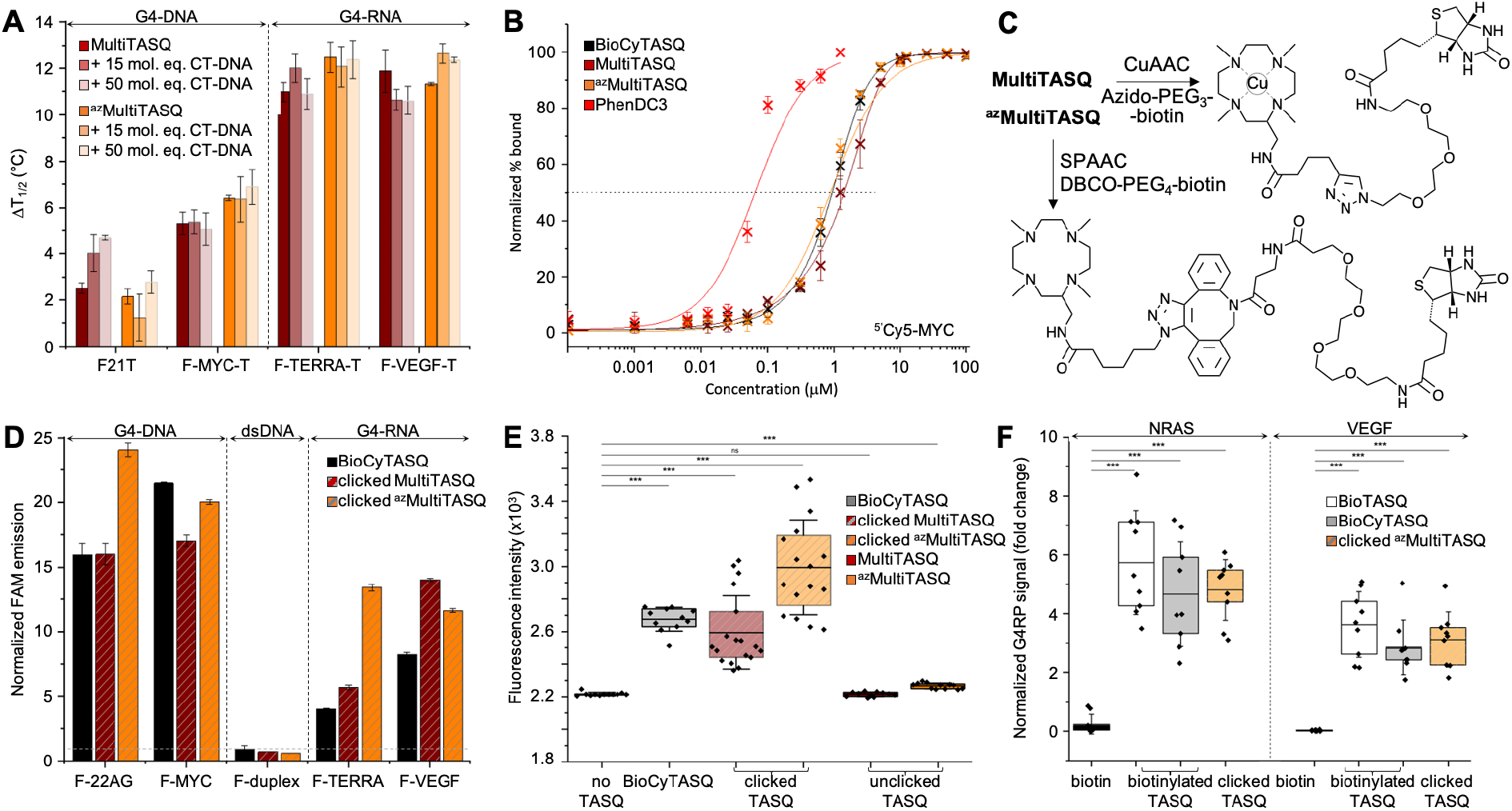
**A**. FRET-melting assay results (ΔT_1/2_, in °C) collected with doubly labelled G4s (0.2 μM; DNA: F-21-T, F-Myc-T; RNA: F-VEGF-T, F-TERRA-T) in the presence of MultiTASQs (1.0 μM) and increasing amounts of unlabeled duplex-DNA (calf thymus DNA, CT-DNA, 15 and 50 mol. equiv.; n >2). **B**. Ligands’ binding to ^5^’Cy5-Myc G4 (20 nM) monitored by Cy5 fluorescence quenching upon addition of increasing amounts (1 pM to 100 μM) of TASQs (and PhenDC3 as control). **C**. Schematic representation of biotinylation of TASQs by bioorthogonal functionalization of MultiTASQ by CuAAC and ^az^MultiTASQ by SPAAC. **D**. Results of the *in vitro* G4 pull-down protocol (n = 3) performed with FAM-labelled oligonucleotides (1.0 μM; DNA: F-22AG, F-Myc; RNA: F-TERRA, F-VEGF; the hairpin F-duplex as control) and clicked MultiTASQs (BioCyTASQ as control) quantified by the increase in fluorescence during the elution step, normalized to the control (without TASQ). **E**. qPCR pull-down results (SYBR green fluorescence intensity; n = 3) for experiments performed with a 97-nt G4-containing DNA sequence (4 μM) without TASQ (control), with clicked MultiTASQs (40 μM , BioCyTASQ as control) or unclicked MultiTASQs (40 μM, control). **F**. G4RP signals of biotin (control) *versus* clicked MultiTASQ (BioTASQ and BioCyTASQ as controls) *via* RT-qPCR quantification of NRAS and VEGFA mRNA levels in MCF7 (n = 3). *p<0.05, **p<0.01, ***p<0.001.

### Click chemo-precipitation of G4s: the fluorescence-based protocol

G4-click-CP was performed with both clicked MultiTASQ and ^az^MultiTASQ (along with BioCyTASQ as a control) against the oligonucleotides used for FRET-melting assay with only the 5’-FAM label (*i.e.*, two DNA G4s: F-22AG and F-Myc, two RNA G4s: F-TERRA and F-VEGF, along with F-duplex as a control, Table S2). MultiTASQ was coupled with azide-PEG_3_-biotin by CuAAC and ^az^MultiTASQ with dibenzocyclooctyne-PEG4-biotin by SPAAC (Figure 2C and the Supp. Info.). MultiTASQ was clicked to azide-PEG_3_-biotin in the presence of an over-stochiometric amount (2.5 mol. equiv.) of copper to take into account the copper chelation by the central cyclen template. The click mixture was prepared in water by mixing (MeCN)_4_Cu·BF_4_ with THPTA ((tris(3-hydroxypropyltriazolylmethyl)amine) before the addition of sodium ascorbate; MultiTASQ was separately mixed with a slight excess or azido-PEG4-Biotin (1.1 mol. equiv.) in a 1:1 mixture of water and 1-butanol (1:1). The two solutions were then mixed and stirred at 25 °C for 1 h (an HPLC-MS monitoring allowed for assessing the efficiency of the CuAAC, if needed). Of note: *i*- the final proportion of 1-butanol is 2% only, which is compatible with the stability of G4s in the condition of the experiments; and *ii*- the demetallation of clicked MultiTASQ, usually performed with Na_2_S treatment,^23, 32^ is avoided here for reproducibility issue related to the loss of material during the precipitation step (the presence of copper within the cyclene template did not affect the properties of the TASQ, Figure S11). ^az^MultiTASQ was mixed with a slight excess of dibenzocyclooctyne (DBCO)-PEG4-biotin conjugate (1.1 mol. equiv.) in water and stirred for 1 h at 37°C (again, an HPLC-MS monitoring allowed for assessing the efficiency of the SPAAC, if needed). Of note: in both instances, we performed affinity control experiments (FRET-melting) that showed that the biotinylation of TASQs by click chemistry does not affect their G4-interacting properties (Figure S11).

The clicked TASQs (10 μM) were then mixed with nucleic acids (1 μM) and streptavidin-coated magnetic beads for 1 h at 25°C before the isolation of DNA (or RNA)/TASQ/beads by magnetic immobilization and release of captured DNA (or RNA) by a thermal denaturation step (8 min at 90 °C). The efficiency of the capture was quantified by the FAM emission calculated after the denaturation step, expressed as fold enrichment after normalization to the control experiment performed without TASQ. As seen in Figure 2D, the three TASQs were found to capture G4s with a similar efficiency; however, contrarily to FRET-melting results, their performance was better with G4-DNA (averaged enrichment = 16.5- and 22.1-fold for clicked MultiTASQ and ^az^MultiTASQ, respectively) than with G4-RNA (averaged enrichment = 9.8- and 12.5-fold for clicked MultiTASQ and ^az^MultiTASQ, respectively). Again, these results were in line with those of BioCyTASQ (averaged enrichment = 18.7- and 6.1-fold for DNA and RNA G4s, respectively) and BioTriazoTASQ.^23^ Quite satisfyingly, none of them were able to pull dsDNA down (between 0.6 and 0.9-fold enrichment with F-duplex), confirming the excellent G4-selectivity of TASQs.

### Click chemo-precipitation of G4s: the qPCR-based protocol

It was thus of interest to assess the G4-capturing ability of TASQ in more biologically relevant conditions. To this end, we included a G4-forming sequence in a 97-nucleotide long DNA strand (Table S3), devoid of fluorescent tags, which makes its detection possible only through qPCR analyses.^33–34^ G4-click-CP was here performed without TASQ (control), with both MultiTASQ and ^az^MultiTASQ (10 μM) either unclicked (controls) or clicked to biotin derivatives, along with BioCyTASQ. As above, MultiTASQ was coupled with azide-PEG_3_-biotin by CuAAC (or not) and ^az^MultiTASQ with dibenzocyclooctyne-PEG4-biotin by SPAAC (or not), and incubated of the G4-containing DNA strand (1 μM) in the presence of the streptavidin-coated magnetic beads for 2 h at 25 °C. After isolation of the DNA/TASQ/beads complexes by magnetic immobilization, the captured DNA was released by a thermal denaturation step (8 min at 90 °C) and quantified through qPCR amplification (expressed as SYBR Green fluorescence intensity, FI). As seen in Figure 2E, no fluorescence increase was observed for the controls (FI = 2215 ± 13 and 2266 ± 18 with unclicked TASQs *versus* 2218 without TASQ; ΔFI = −3 and 48, respectively), thus confirming the need of a biotin bait for isolating G4s. Both clicked MultiTASQ and ^az^MultiTASQ efficiently pulled G4 down (FI = 2594 ± 225 and 2924 ± 291, respectively; ΔFI = 376 and 706), in a manner that is reminiscent to what is observed with BioCyTASQ (FI = 2675 ± 73; ΔFI = 457). These results thus confirmed that of the fluorescence-based G4-click-CP.

## IV. Cell-based applications of MultiTASQs

### G4RP protocol with clicked ^az^MultiTASQ

The two aforementioned G4-click-CP protocols were purely *in vitro* manipulations. To go a step towards using TASQ baits in more relevant conditions, we considered both the G4RP protocol and *in situ* click imaging. We implemented ^az^MultiTASQ for the former and MultiTASQ for the latter.

The G4-RNA precipitation (G4RP) protocol was developed to detect folded G4s *in vivo*.^20-21, 24^ G4RP hinges on the cross-linking of naturally occurring G4s in living cells using formaldehyde prior to isolating them from cell lysates by affinity precipitation with BioTASQ. The G4RP protocol was validated by RT-qPCR analysis against well-established G4-containing transcripts including VEGF (see above) and a sequence found in the UTR region of the NRAS mRNA. To date, the G4RP-RT-qPCR protocol was performed with BioTASQ only;^20-21, 24^ we thus decided to evaluate the properties of BioCyTASQ and a clicked ^az^MultiTASQ and to compare them with the initially used TASQ bait.

The interaction between TASQ and VEGF has already been investigated above; we thus checked their binding to NRAS by FRET-melting, performing the experiments with F-NRAS-T (0.2 μM) and TASQs (1 μM, 5 mol. equiv.) in the absence or presence of an excess of competitive CT-DNA (15 or 50 mol. equiv.). The results seen in Figure S11A indicated that NRAS is efficiently stabilized by the TASQs with ΔT_1/2_ = 11.8 ± 0.7 and 14.6 ± 1.1 °C for MultiTASQ, ^az^MultiTASQ, respectively, with an exquisite selectivity (^FRET^S > 0.96). We then checked that clicked TASQs efficiently pulled F-NRAS down *via* fluorescence G4-click-CP: the enrichment seen in Figure S11B (5.5 ± 0.8 and 2.4 ± 0.3 for clicked MultiTASQ and ^az^MultiTASQ, respectively, *versus* 5.2 ± 0.9 for BioCyTASQ) confirmed that TASQs are indeed valuable baits for isolating this transcript *in vitro*.

The G4RP results depicted in Figure 2F confirmed that the sterically demanding DBCO-based linker (comprising 1 triazole, 2 phenyls and 1 azacyclooctane, Figure 2C) of clicked ^az^MultiTASQ does not hamper proper interaction with G4s *in vivo*. Indeed, MCF7 cells were trypsinyzed and then cross-linked with formaldehyde for 5 min prior to be resuspended in G4RP buffer and lysed (mechanical disruption). The lysate was then incubated with biotinylated (BioTASQ, BioCyTASQ) or pre-clicked ^az^MultiTASQs (along with biotin as control, 100 μM) in the presence of magnetic beads for 1 h at 4 °C. The beads were then isolated (magnetic immobilization), washed with G4RP buffer and subjected to a thermal treatment (70 °C for 1 h) to both reverse the cross-link and free the captured nucleic acids. RNA fragments were isolated thanks to TRIZOL extraction and the quantity of NRAS and VEGF transcripts assessed by RT-qPCR. In these conditions, clicked ^az^MultiTASQ efficiently enriched both NRAS and VEGF transcripts (enrichment = 4.8 ± 1.0 and 3.1 ± 0.9, respectively), less efficiently than BioTASQ (5.7 ± 1.7 and 3.6 ± 1.1, respectively) but with a better reproducibility, and more efficiently than BioCyTASQ (4.6 ± 1.8 and 2.8 ± 0.9, respectively). When compared to the biotin control (enrichment = 0.2 ± 0.3 and 0.02 ± 0.02, respectively), the two mRNA transcripts are enriched *ca*. >20-and >100-fold by the TASQs.

### Click imaging with MultiTASQ

To further exploit the versatility of MultiTASQs, we used their clickable handle *in situ* click imaging protocols to image G4 landscapes by clicking TASQs, once in their cellular G4 sites, with fluorescent partners. This approach is different from the pre-targeted G4 imaging we previously reported on,^22–23^ as the very nature of the clickable appendages of MultiTASQs (small size, no H-bonding ability) ensures that the target engagement of TASQs in cells is not biased, as it could be the case with the biotin appendage. To this end, we adapted the *in situ* click imaging protocol initially developed with PDS-α:^13^ MCF7 human breast cancer cells were incubated either live (10 μM, 24 h) or after fixation (20 μM, 1 h) with MultiTASQ (to demonstrate the modularity of this approach). Bioorthogonal click reactions were then performed in cells with either AF488-azide or AF594-azide (to further demonstrate its modularity) by CuAAC. Nuclei were subsequently stained with DAPI and images were collected by confocal laser scanning microscopy. The pattern seen in Figure 3 corresponds to what has been described for the twice-as-smart probe N-TASQ (direct labelling)^35–36^ and biotinylated TASQ (pre-targeted imaging):^22–23^ a dense nucleolar staining (yellow arrow) along with smaller, discrete nucleoplasmic *foci* (white arrow) ascribed to G4 clusters that fold during DNA transactions as they localize where DNA is at work (no DAPI staining). This approach, though qualitative in nature, provides high-quality images (high brightness, signal-to-noise ratio and *foci* definition) that are currently being exploited to assess whether and how G4-interacting agents (stabilizers^37^ or destabilizers)^34, 38^ modulate G4 landscapes in cells. These results will be reported in due course.

**Figure 3.**
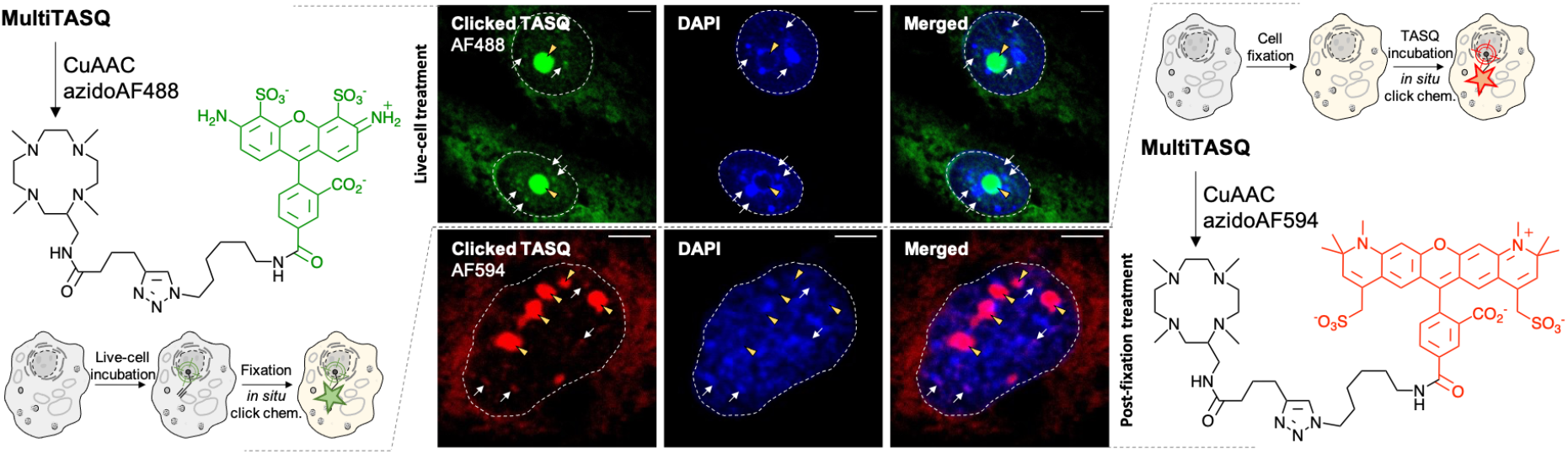
Schematic representation of *in situ* click G4 imaging protocols performed with MultiTASQ and either AF488-azide (left) or AF594-azide (right). Central panel: representative confocal images of MCF7 cells collected through DAPI (<425 nm), GFP (500-550 nm) and Cy3 (>580 nm) filters. Cells are either treated live with MultiTASQ (10 μM, 24 h) prior to be fixed (upper panel) or fixed prior to be treated with MultiTASQ (20 μM, 1 h) (lower panel) and then subjected to Cu-catalyzed bioorthogonal labelling with either AF488-azide (upper panel) or AF594-azide (lower panel). Yellow arrows indicate nucleoli; white arrows indicated clicked MultiTASQ accumulation sites, ascribed to G4 *foci*; dashed lines delineate cell nuclei.

## Conclusion

In conclusion, the design of MultiTASQs makes them benefit from the very high versatility and wide scope of click chemistry. The presence of azide/alkyne appendages offers more experimental opportunities than that the biotinylated TASQs can offer as these appendages do not create unwarranted interactions *i*- within the structure of the TASQ itself (avoiding internal structural poisoning), and *ii*- with cellular components (preventing ligand dispersal), both mainly occurring *via* H-bond formation. Because both the biological activity and the target engagement of the TASQs are not biased, reliable live-cell experiments are possible.

We applied here in the G4 field only a minor fraction of the many possibilities offered by click chemistry.^39–40^ However, the applications we developed with the clickable TASQs say much about how they are likely to become important companions for deciphering G4 biology. First, the click chemo-precipitation of G4s (G4-click-CP) helps assess whether suspected G4-forming sequences (both DNA and RNA) do actually fold into G4 structures in cells (G4RP^20^ and G4DP^41^ protocols, respectively). It also helps determine the extent to which G4 landscapes is modulated in cells by external effectors (*e.g.*, ligands) in a quantitative manner. The modularity of these protocols, which are implementable in either targeted (RT-qPCR analysis) or genome-wide manner (sequencing), makes them fully complementary to existing methods (for DNA G4s: G4-seq^42^ using the G4 ligand PDS,^12^ or G4 ChIP-seq,^43^ BG4-ChIP-seq^44^ and G4P ChIP-seq^45^ using the G4-sepcific antibody BG4;^46^ for RNA G4s: rG4-seq^47^ using PDS, or BG4-RNA-IP^48^ and rG4IP^49^ using BG4), which altogether constitute a unique array of techniques to accurately portray G4 biology. Second, the *in situ* click imaging of G4s provides a more qualitative but also straightforward way to track G4s in cells. This fluorescence microscopy technique does not provide fine details about the sequences involved but a unique and dynamic visualization of the cellular G4 sites, which could be amenable to mechanistic interpretation *via* both colocalization studies (with organelle markers, DNA damage markers, etc.) and longitudinal studies (*e.g*., upon ligand treatment). Optical imaging techniques aiming at tracking G4 which can also be more quantitative in nature and implementable as high-throughput screens such as the flow cytometry-based protocol BG-flow,^50^ based on the BG4, used to discriminate between cell status and monitor ligand-mediated modulations of G4 signatures.

Clickable TASQs thus allow for the implementation of protocols that represent two faces only of an unicum of techniques devoted to uncovering the finest details of G4 biology. With this growing portfolio of tools in hands, new experimental, strategic and mechanistic opportunities are now available for pushing the G4 field ever further.

## Material and Methods

### 1. Chemistry & oligonucleotides

The synthesis and chemical characterization of MultiTASQ and ^az^MultiTASQ are described in the Supporting Information; the oligonucleotides used in this study along with the preparation of their higher-order structures are also described in the Supporting Information. The click protocols are the following:

#### 1a. CuAAC

5 μL of a 1-M solution (DMSO) of (MeCN)_4_Cu·PF_6_ in DMSO were mixed with 7 μL of a 1-M solution (H_2_O) of THPTA (tris(3-hydroxypropyltriazolylmethyl)amine), the mixture rapdidly turned to a dark blue solution (Cu(II) salt). To this solution were added 10 μL of 1-M solution (H_2_O) of sodium ascorbate to provide a colorless solution of Cu(I) salt. Separately, 20 μL of a 5-mM solution (water/1-Butanol 1:1) of MultiTASQ were mixed with 1.1 μL of a 100-mM solution (H_2_O) of Azido-PEG4-Biotin conjugate, to which 4μL of the aforementioned Cu(I) solution were added. The reaction was stirred for 1 h at 25°C (HPLC-MS monitoring) and the stirring was stopped for a blue precipitate to form (Cu(II) salt). After centrifugation, the supernatant was removed and the clicked MultiTASQ used without further purification.

#### 1b. SPAAC

20 μL of a 1-mM solution (H_2_O) of ^az^MultiTASQ were mixed with 2.2 μL of a 10-mM solution (H_2_O) of dibenzocyclooctyne-PEG4-biotin conjugate (DBCO-PEG4-Biotin). The reaction was stirred for 1 h at 37°C for 1 hour (HPLC-MS monitoring) after which the clicked ^az^MultiTASQ was used without further purification.

### 2. Affinity measurements

#### 2a. FRET-melting assay

FRET-melting experiments were performed in a 96-well format using a Mx3005P qPCR machine (Agilent) equipped with FAM filters (λ_ex_ = 492 nm; λ_em_ = 516 nm) in 100 μL (final volume) of CacoK10 (Table S4) for F21T or CacoK1 (Table S4) for F-Myc-T, F-Terra-T and F-VEGF-T, with 0.2 μM of labelled oligonucleotide (Table S1) and 1 μM of TASQ. Competitive experiments were performed with labelled oligonucleotide (0.2 μM), 1 μM TASQ and either 15 (3 μM)) or 50 mol. equiv. (10 μM) of the unlabeled competitor calf thymus DNA (CT-DNA). After an initial equilibration step (25°C, 30 s), a stepwise increase of 1°C every 30s for 65 cycles to reach 90°C was performed, and measurements were made after each cycle. Final data were analyzed with Excel (Microsoft Corp.) and OriginPro®9.1 (OriginLab Corp.). The emission of FAM was normalized (0 to 1), and T_1/2_ was defined as the temperature for which the normalized emission is 0.5; ΔT_1/2_ values are means of 3 triplicates.

#### 2b. Apparent K_D_ measurement

To a solution of Cy5-myc (20 nM) in 50mM TrisHCl/Triton buffer (Table S4) was added various concentrations (from 100 μM to 1 pM) of TASQs (and PhenDC3 as control). After mixing the solutions for 1 hour at 25°C, the fluorescence was measured using a ClarioStar® machine (BMG Labtech) equipped with Cy5 filter (λ_ex_ = 610 nm; λ_em_ = 675 nm). Data were analyzed with Excel (Microsoft Corp.) and OriginPro^®^9.1 (OriginLab Corp.); the values were normalized ranging from 0 to 100, and the percentage of ligand bound was calculated subtracting the normalized values to 100. Three biological triplicates (n = 3) were used.

### 3. G4-click-CP

#### 3a. Fluorescence pull-down assay

The streptavidin MagneSphere^®^ beads (Promega) were washed 3 times with 20 mM TrisHCl/MgCl_2_ buffer (Table S4). TASQs (10 μM) was mixed with 5’-labelled oligonucleotides (F-ON, 1 μM: F-22AG, F-Myc, F-duplex, F-TERRA and F-VEGF, Table S2), MagneSphere^®^ beads (32 μg) in the same TrisHCl buffer (320 μL final volume) and stirred for 1 h at 25 °C. The beads were immobilized (fast centrifugation (< 2 s), magnet) and the supernatant removed. The solid residue was resuspended in 320 μL of TBS 1X buffer, heated for 8 min at 90 °C (gentle stirring 800 r.p.m.) and then centrifuged for 2 min at 8900 rpm. The supernatant was taken up for analysis (magnet immobilization), after being distributed in 3 wells (100 μL each) of a 96-well plate, using a ClarioStar^®^ machine (BMG Labtech) equipped with FAM filters (λ_ex_ = 492 nm; λ_em_ = 516 nm). Data were analyzed with Excel (Microsoft Corp.) and OriginPro^®^9.1 (OriginLab Corp.); normalized FAM emission values are means of 3 triplicates; each analysis comprises: a/ 3 control wells with F-ON and beads only (in order to quantify the non-specific F-ON/bead binding, the FAM emission of the solution was normalized to 1); and b/ 3 wells comprising solutions that resulted from experiments performed with F-ON, TASQ and beads (in order to quantify the capture capability of TASQ when compared to the control experiments).

#### 3b. qPCR pull-down assay

The pull-down experiments were performed as above (*cf*. 3a), with the following modifications: a/ the oligonucleotide used was changed for a 97-nt DNA (Table S3) described in Jamroskovic *et al*.^33^ and adapted in Mitteaux *et al*.^34^ at the center of which the G4-forming sequence d[(GGGCA)4] is included; b/ the buffer was replaced by the G4RP buffer (Table S4); c/ the incubation time was changed for 2 h at 25 °C; d/ the output was changed for qPCR analyses: polymerase reactions were carried out in triplicate in 96-well format using a Mx3005P qPCR machine (Agilent) equipped with FAM filters (λ_ex_ = 492 nm; λ_em_ = 516 nm) in 20 μL (final volume) of G4-1R primer (1 μL, 300 nM), TASQ/ODN mixture (3.7 μL) in 10 μL iTaq™ Universal SYBR^®^ Green Supermix (Bio-Rad) + KCl (5.3 μL, 100 mM). After a first denaturation step (95 °C, 5 min), a two-step qPCR comprising a hybridization step (85 °C, 10 s) and an elongation step (60 °C, 15 s) for 33 cycles was performed, and measurements were made after each cycle. Final data were analyzed with OriginPro^®^9.1 (OriginLab Corp.). The starting emission (1^st^ qPCR cycle) of SYBR Green (FI) was set to 2200 and the FI at the 33th cycle was used for calculation. Three biological triplicates (n = 3) were used.

### 4. Cell-based investigations

#### 4a. G4RP-RT-qPCR protocol

MCF7 cells were seeded at 7M cells per 175 cm^2^ flask. After overnight adhesion, the medium was changed and cells were further cultured for 48 h before being trypsinized and then crosslinked using 1% (v/v) formaldehyde in fixing buffer (Table S4) for 5 min at 25 °C. The crosslink was then quenched with 0.125 M glycine for 5 min and washed and rinsed (with DEPC-PBS, Table S4). Cells were resuspended in G4RP buffer + 0.1% (v/v) SDS and then manually disrupted (syringe). After centrifugation (13 200 rpm, 10 min), the collected lysates (5% of which were collected as input control) were incubated with 100 μM TASQs (or 100 μM biotin as control) and 90 μg of MagneSphere^®^ beads (Promega) for 2 h at 4°C. Magnetic beads were then washed (5 min, twice), before being resuspended in DEPC-PBS buffer supplemented with 0.4 U RNAse OUT. The beads were then incubated at 70 °C for 2 h to release captured G4-forming targets from the beads as unfolded. TRIZOL (1 mL) was then used to extract the RNA (using manufacturer’s instructions) and cleaned with RNA Clean-up protocol (using manufacturer’s instructions) at 25 °C. The primer sets used for RT-qPCR are NRAS-forward and NRAS-reverse, and VEGF-forward and VEGF-reverse (Table S3). Extracted RNA was reverse transcribed with Superscript III (InvitrogenTM 18080-044) and random hexamer primers (InvitrogenTM N8080127) using manufacturer’s protocol to generate cDNA. cDNAs were quantified for target mRNAs using iTaqTM Universal 2X SYBR^®^ Green Supermix (Bio-Rad) and specific primer sets with three technical replicates in each assay. C(t) values of pull-down samples were normalized to the input control. Three biological replicates were used for all RT-qPCR-based quantifications. Final data were analyzed with Excel (Microsoft Corp.) and OriginPro^®^9.1 (OriginLab Corp.). For statistical hypothesis student’s *t*-test and Welch’s unequal variances *t*-test were used depending on variances equality.

#### 4b. *in situ* click imaging

MCF7 cells were seeded on glass coverslips in a 24 well-plate for 24 h at 37 °C. Cells were either treated live with MultiTASQ (10 μM in DMEM, 24 h) then fixed (ice cold methanol, 10 min), or fixed and treated with MultiTASQ (20 μM in PBS, 1 h). Coverslips were washed with PBS (5 min, thrice), and click staining performed with AF488- or AF594-azide (1 μM) in PBS containing 0.05% IGEPAL CA-630, 1 mM CuSO4 and 10 mM sodium ascorbate for 30 min, washed with PBS+0.1% Triton (5 min, thrice), incubated with DAPI (10 min, 1 μg/mL in PBS) and mounted with Fluoromount. Confocal imaging was performed using a confocal laser-scanning microscope (Leica TCS SP8) with a 63× objective lens and LASX software (Leica Microsystems CMS GmbH). Image processing was carried out using ImageJ.

## Supporting Information

Synthesis and characterizations of MultiTASQ (Figures S1-S5) and ^az^MultiTASQ (Figures S6-S10); oligonucleotide sequences (Tables S1-S3) and preparation; additional results for FRET-melting experiments and pull-down assays (Figure S11).

## Acknowledgments

This work was supported by the CNRS (I.E.V. and D.M.), iSITE BFC (COMUE UBFC, PIA2, grant n° UB21018.MUB.IS, for D.M.), and the European Union (PO FEDER-FSE Bourgogne 2014/2020 programs, grant n° BG0021532 for D.M.). We also thank C. Deulvot & E. Noirot, DImaCell, AgroSup Dijon, INRAe, UBFC Dijon for technical assistance regarding confocal microscopy.

