## Supplementary Material for "The multivalent G-quadruplex (G4)-ligands MultiTASQs allow for versatile click chemistry-based investigations"

ICMUB, CNRS UMR6302, UBFC Dijon, 9 Avenue Alain Savary, 21078 Dijon, France

**- - Supporting Information - -**

**I. Chemistry**······································page S01
Synthesis of MultiTASQ······························page S03
Synthesis of $^{az}$MultiTASQ······························page S09
**II. Oligonucleotides**······································page S16
**III. Additional *in vitro* results**······························page S17

### I. Chemistry

**Ia. Generalities.** TSTU was purchased from Iris Biotech GmbH. Solvents were purchased from VWR or Carlo Erba. All other chemicals were from Sigma Aldrich, Acros organics, or Fisher Scientific. Cyclen was a gift from Chematech (Dijon, France). Media and supplements for cell culture were bought from Dutscher SAS. Unless noted otherwise, all commercially available reagents and solvents were used without further purification. Dicalite was purchased from Carlo Erba. TLC were carried out on Merck DC Kieselgel 60 F-254 aluminum sheets. The spots were visualized *via* staining with ninhydrin or through illumination with a UV lamp ($\lambda$ = 254 nm). Column chromatography purifications were performed manually on silica gel (40-63 $\mu$m) from Sigma-Aldrich (technical grade). Dry $CH_2Cl_2$ (HPLC-grade) was dried over alumina cartridges using a solvent purification system PureSolv PS-MD-5 model from Innovative Technology. HPLC-gradient grade $CH_3CN$ used for HPLC-MS analyses was obtained from Carlo Erba. $CH_3CN$ used in semi-preparative RP-HPLC purifications was obtained from VWR (technical, +99% but distilled prior to use). All aq. mobile-phases for HPLC were prepared using water purified with a PURELAB Ultra system from ELGA (purified to 18.2 MΩ.cm). Yields were calculated based on isolation of the compounds.

**Ib. Instruments and methods.** Lyophilization was performed with a Christ Alpha 2-4 LD plus. $^{1}H$-, and $^{13}C$- NMR spectra were recorded on Bruker spectrometers, an Avance Neo 500 MHz equipped with a 5 mm BBOF iProbe and Avance III HD.Chemical shifts are expressed in parts per million (ppm) from the residual non-deuterated solvent signal summarized in 2010 by Fulmer *et al*. (*Organometallics* **2010**, *29*, 2176-2179) *J* values are expressed in Hz. Coupling constants (*J*) are reported in hertz (Hz). Standard abbreviations indicating multiplicity were used as follows: s = singlet, d = doublet, t = triplet, q = quadruplet, m = multiplet, br = broad. High-resolution mass spectrometry analyses were recorded on a LTQ Orbitrap XL mass spectrometer (Thermo Scientific) equipped with an electrospray ionization source (HESI 2). The following source parameters were used if no further specification is mentioned: Heater Temperature: 50°C, Gas Flow: Sheath 15 / Aux 10 / Sweep 0 , Spray Voltage: 4 kV, Capillary Temperature: 275°C, Capillary Voltage: 22 V, Resolution (m/z = 400): 60 000. HPLC-MS analyses were performed on a Thermo-Dionex Ultimate 3000 instrument (pump + autosampler at 20 °C + column oven at 25 °C) equipped with a diode array detector (Thermo-Dionex DAD 3000-RS) and a MSQ Plus single quadrupole mass spectrometer. The corresponding low-resolution mass spectra (LRMS) were recorded with this latter mass spectrometer, with an electrospray (ESI) source (HPLC-MS coupling mode) HPLC systems were equipped with a Phenomenex Kinetex C18 column, 2.6µm, 2.1 × 50 mm or a Jupiter Proteo 4 µm 90Å column, 250 x 4.6 mm. Two analytical methods: 1/ *Method A*: from 5% to 100% MeCN/$H_2O$+0.1% formic acid (FA) in 7 min; 2/ *Method B*: from 5% to 15% MeCN/$H_2O$+0.1% FA in 5 min, from 15% to 70% MeCN/ $H_2O$+0.1% FA in 20 min, from 70% to 100% MeCN/ $H_2O$+0.1% FA in 3 min. Purifications by semi-preparative HPLC were performed on a Thermo-Dionex Ultimate 3000 instrument equipped with a RS Variable Detector (four distinct wavelengths). HPLC system was equipped with a Jupiter Proteo 4 µm 90Å column (250 x 21.2 mm, AXIA packed). Final compounds were analyzed with an ACCUCORE column, Method C: stationary at 0% for 0.6 min, from 0% to 20% MeCN/H2O + 0.1% formic acid (FA) in 2.4 min and then from 20% to 100% MeCN/H2O + 0.1% formic acid (FA) in 2.6 min, the gradient stays stationary for 1.5 min at 100% MeCN, the run is stopped at 11 min.

**Ic. Synthesis and characterizations.**

**MultiTASQ synthesis – Step 1:**

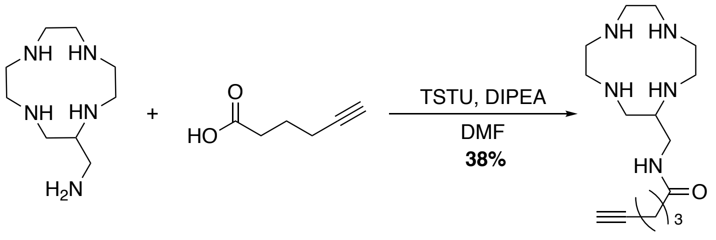


To a solution of hexynoic acid (167 µL, 0.61 mmol, 0.9 equiv.) and DIPEA (315 µL, 1.34 mmol, 2 equiv.) in DMF (2 mL) was added TSTU (228 mg, 0.74 mmol, 1.1 equiv.) and the solution was allowed to stir 1 hour until complete conversion of the starting material assessed through HPLC-MS. To a solution of aminomethylcyclen (AMC, 134,7 mg, 0.67 mmol, 1 equiv.) in DMF (10 mL) at room temperature was added dropwise the previously prepared reaction mixture containing N-hydroxylsuccimide hex-5-ynoate (1 mL/3 hrs). The reaction was carefully monitored by HPLC-MS. Upon completion, the solution was concentrated under vacuum, the residue diluted in water and purified by SemiPrep-HPLC in a $H_2O/CH_3CN$ + 0.1% TFA mixture (gradient of 5 to 50% over 35 minutes, column *B*, r.t: 12.5 minutes). After evaporation of the solvents, the alkynylated AMC was obtained (187.4 mg, 0.25 mmol, 38% yield). $^1$H NMR: (500 MHz, $D_2O$): δ. 3.70-2.77 (m, 14H), 2.37 (t, J = 7.4 Hz, 1H), 2.23 (m, 2H), 1.80 (m, 1H), 1.36 (m, 6H). $^{13}$C NMR: (126 MHz, $D_2O$): δ. 177.04, 162.83, 119.09, 117.16, 115.23, 84.58, 69.91, 52.26, 46.42, 44.18, 43.89, 42.08, 40.51, 39.01, 23.75. ESI-HRMS: $[M+H]^+$ m/z = 296.243 (calcd. for $C_{15}H_{30}N_5O$: 296.245). HPLC-MS characterization (Phenomenex Kinetex C18 column, 2.6µm, 2.1 × 50 mm) *via Method A* (from 5% to 100% MeCN/$H_2O$+0.1% formic acid (FA) in 7 min): retention time = 0.377 min; purity: >92% at 201 nm, >90% at 214 nm; m/z = 296.4 $[M+H]^+$ (cf. Figure S1).

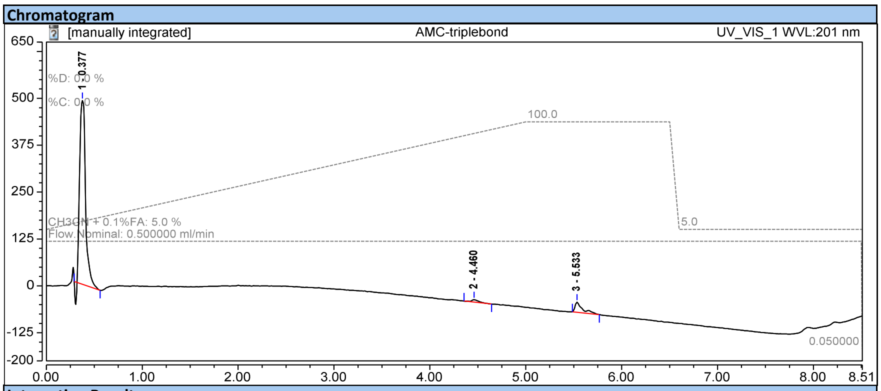


**Integration Results**

| No. | Peak Name | Retention Time min | Area mAU*min | Height mAU | Relative Area % | Relative Height % | Amount n.a. |
|---|---|---|---|---|---|---|---|
| 1 | | 0.377 | 34.340 | 489.558 | 92.08 | 93.66 | n.a. |
| 2 | | 4.460 | 0.542 | 6.308 | 1.45 | 1.21 | n.a. |
| 3 | | 5.533 | 2.413 | 26.849 | 6.47 | 5.14 | n.a. |
| **Total:** | | | **37.295** | **522.715** | **100.00** | **100.00** | |

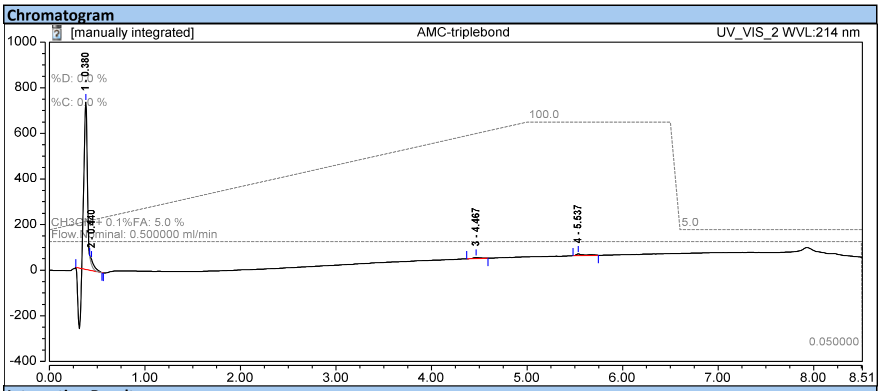


**Integration Results**

| No. | Peak Name | Retention Time min | Area mAU*min | Height mAU | Relative Area % | Relative Height % | Amount n.a. |
|---|---|---|---|---|---|---|---|
| 1 | | 0.380 | 23.404 | 733.669 | 90.02 | 95.41 | n.a. |
| 2 | | 0.440 | 1.397 | 22.710 | 5.37 | 2.95 | n.a. |
| 3 | | 4.467 | 0.397 | 4.912 | 1.53 | 0.64 | n.a. |
| 4 | | 5.537 | 0.801 | 7.692 | 3.08 | 1.00 | n.a. |
| **Total:** | | | **25.998** | **768.984** | **100.00** | **100.00** | |

**Figure S1**. HPLC profiles of the alkynylated AMC

### MultiTASQ synthesis – Step 2:

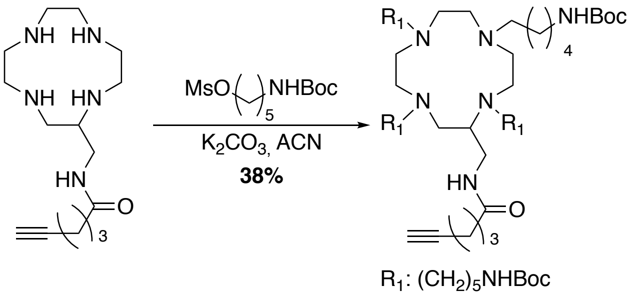


To a solution of alkynylated AMC (552.6 mg, 0.73 mmol, 1 equiv.) in $CH_3CN$ (7 mL) was added 5-((tert-butoxycarbonyl)amino)pentyl methanesulfonate (1654.1 mg, 5.9 mmol, 8 equiv.) and $K_2CO_3$ (814.2 mg, 5.9 mmol, 8 equiv.). The solution was stirred at 70°C for 72 hours, being carefully monitored by HPLC-MS. Upon completion, the crude mixture was filtered, concentrated under vacuum and diluted in water prior to be purified by SemiPrep-HPLC in a $H_2O$/MeCN + 0.1% TFA mixture (gradient of 5 to 60 % over 50 minutes, column *A*, r.t: 22 minutes). The Boc-protected compound was obtained after removal of the solvents by lyophilization (287.4 mg, 0.28, mmol, 38 % yield). $^1$H NMR: (500 MHz, Methanol-$d_4$) δ 4.28 (t, $J$ = 6.2 Hz, 1H), 3.67 (s, 1H), 3.17-3.03 (m, 15H), 2.98-2.77 (m, 10H), 2.74-2.69 (m, 3H), 2.35 (t, $J$ = 7.5 Hz, 2H), 2.29 (s, 1H), 2.24 (m, 2H), 2.05 (s, 1H), 1.86-1.78 (m, 7H), 1.77-1.65 (m, 7H), 1.61-1.50 (m, 14H), 1.48-1.36 (m, 36H). ESI-HRMS: $[M+H]^+$ m/z = 1036.811 (calcd. for $C_{55}H_{105}N_9O_9$: 1036.810). HPLC-MS characterization (*Method A*): retention time = 4.06 min; purity: >98% at 201 nm, >95% at 214 nm; m/z = 1037.3 $[M+H]^+$ (cf. Figure S2).

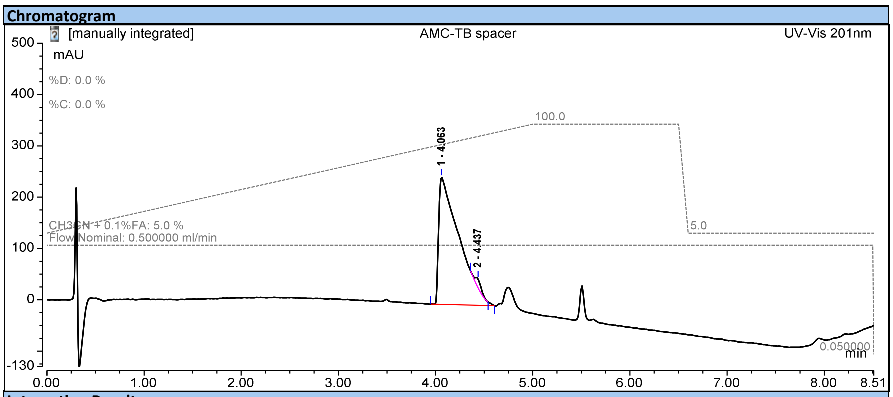


**Integration Results**

| No. | Peak Name | Retention Time min | Area mAU*min | Height mAU | Relative Area % | Relative Height % | Amount n.a. |
|---|---|---|---|---|---|---|---|
| 1 | | 4.063 | 60.227 | 247.336 | 98.05 | 93.48 | n.a. |
| 2 | | 4.437 | 1.196 | 17.259 | 1.95 | 6.52 | n.a. |
| **Total:** | | | **61.422** | **264.595** | **100.00** | **100.00** | |

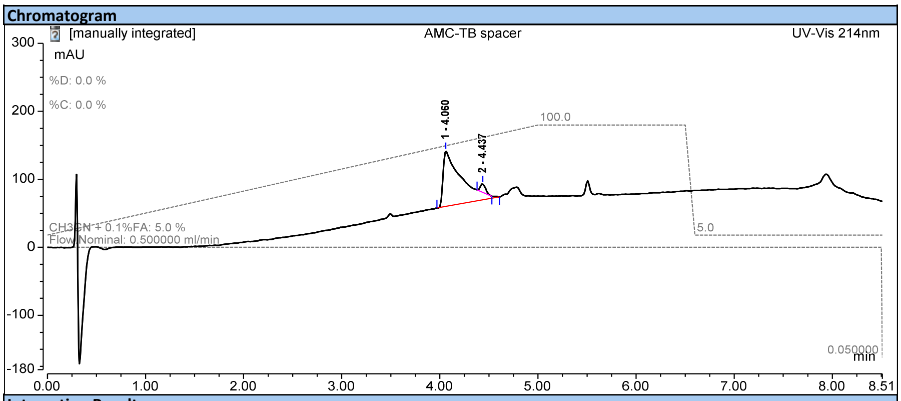


**Integration Results**

| No. | Peak Name | Retention Time min | Area mAU*min | Height mAU | Relative Area % | Relative Height % | Amount n.a. |
|---|---|---|---|---|---|---|---|
| 1 | | 4.060 | 17.179 | 81.628 | 95.71 | 86.90 | n.a. |
| 2 | | 4.437 | 0.770 | 12.300 | 4.29 | 13.10 | n.a. |
| **Total:** | | | **17.949** | **93.928** | **100.00** | **100.00** | |

**Figure S2**. HPLC profiles of the Boc-protected compound

**MultiTASQ synthesis – Steps 3 and 4:**

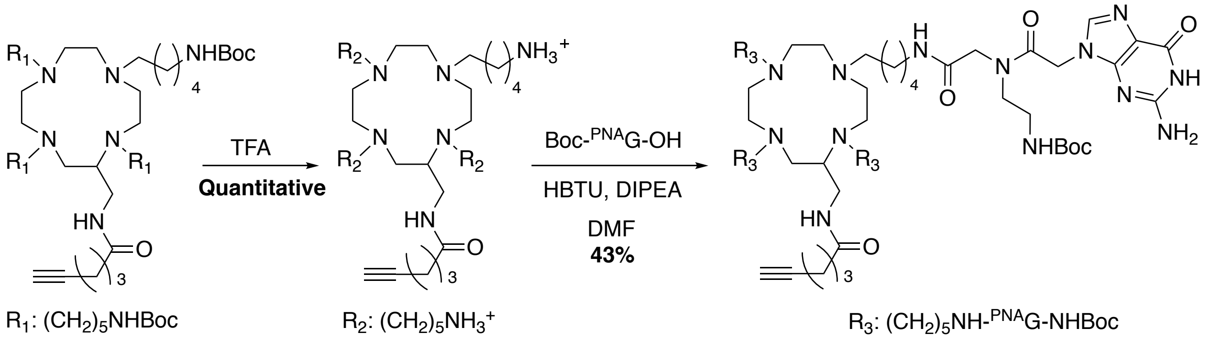


The previously Boc-protected compound (14.8 mg, 0.014 mmol, 1 equiv.) was stirred in TFA (500 μL) for 1 hour. After evaporation of the TFA, the deprotected compound was directly engaged in following reaction without purifications: Boc-$^{PNA}$G-OH (25.8 mg, 0.063 mmol, 4.4 equiv.) and HBTU (23.9 mg, 0.063 mmol, 4.4 equiv.) were dissolved in DMF (1 mL) and DIPEA was added (12 μL, 4 equiv.). This mixture was added to a solution of the previously prepared deprotected compound (14.8 mg, 0.014 mmol, 1 equiv.) and DIPEA (3 μL, 1 equiv.) in DMF (1 mL). The mixture was stirred at 25 °C for 3 hours, being carefully monitored by HPLC-MS. Upon completion, the solution was then concentrated under vacuum, the residue diluted in a mixture of water and MeCN (50/50, 2mL) and purified by SemiPrep-HPLC in a $H_2O$/MeCN + 0.1% TFA mixture (gradient of 5 to 15% over 5 minutes then from 15 to 65% over 50 minutes, column *A*, r.t: 37 minutes). The protected MultiTASQ was obtained after evaporation of the solvents by lyophilization (13.6 mg, 0.006 mmol, 43% yield). $^1$H NMR: (500 MHz, Methanol-$d_4$) δ 8.22 (m, 2H), 7.96 (m, 4H), 5.06 (m, 10H), 4.95 (m, 4H), 4.15 (s, 2H), 3.91 (s, 8H), 3.55 (s, 8H), 3.45-3.25 (m, 14H), 3.16-2.77 (m, 24H), 2.34 (s, 3H), 2.18 (s, 1H), 2.13 (m, 2H), 1.70 (m, 4H), 1.45-1.33 (m, 42H) 1.31 (s, 10H), 1.26-0.94 (m, 14H). ESI-HRMS: $[M+H]^{2+}$ m/z = 1101.130 (calcd. for $C_{99}H_{157}N_{37}O_{21}$: 1101.125). HPLC-MS characterization (*Method A*): retention time = 3.657 min; purity: >96% at 201 nm, >85% at 214 nm, >87% at 280 nm; m/z = 1102.2 $[M+H]^{2+}$ (cf. Figure S3).

**Chromatogram**

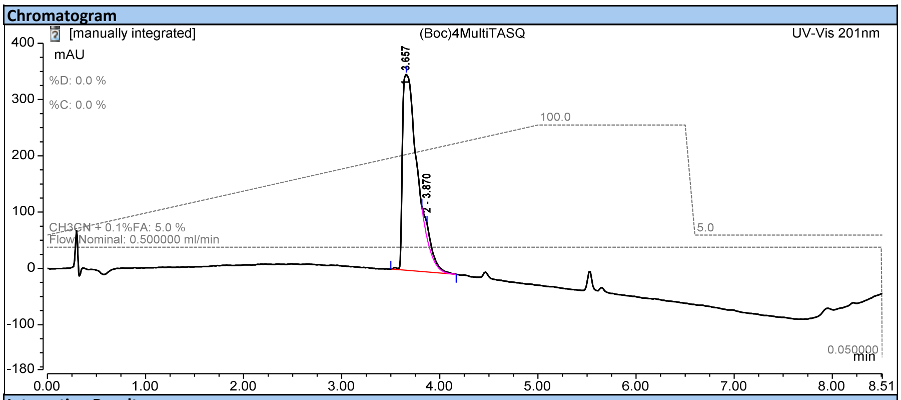


**Integration Results**

| No. | Peak Name | Retention Time<br>min | Area<br>mAU*min | Height<br>mAU | Relative Area<br>% | Relative Height<br>% | Amount<br>n.a. |
|---|---|---|---|---|---|---|---|
| 1 | | 3.657 | 61.814 | 347.755 | 96.28 | 93.33 | n.a. |
| 2 | | 3.870 | 2.388 | 24.844 | 3.72 | 6.67 | n.a. |
| **Total:** | | | **64.201** | **372.598** | **100.00** | **100.00** | |

**Chromatogram**

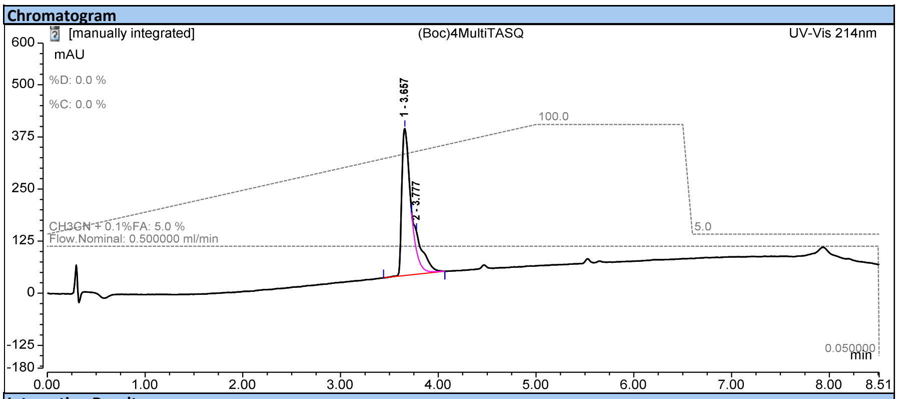


**Integration Results**

| No. | Peak Name | Retention Time<br>min | Area<br>mAU*min | Height<br>mAU | Relative Area<br>% | Relative Height<br>% | Amount<br>n.a. |
|---|---|---|---|---|---|---|---|
| 1 | | 3.657 | 37.997 | 352.648 | 84.79 | 88.58 | n.a. |
| 2 | | 3.777 | 6.818 | 45.447 | 15.21 | 11.42 | n.a. |
| **Total:** | | | **44.815** | **398.095** | **100.00** | **100.00** | |

**Chromatogram**

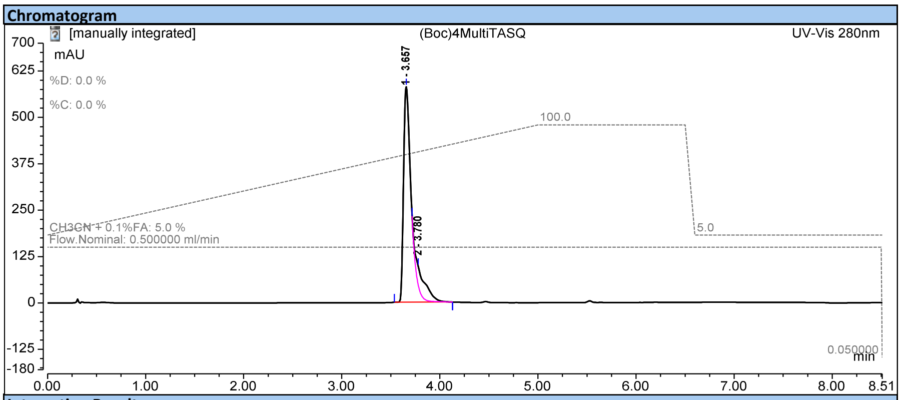


**Integration Results**

| No. | Peak Name | Retention Time<br>min | Area<br>mAU*min | Height<br>mAU | Relative Area<br>% | Relative Height<br>% | Amount<br>n.a. |
|---|---|---|---|---|---|---|---|
| 1 | | 3.657 | 52.264 | 580.342 | 87.07 | 91.98 | n.a. |
| 2 | | 3.780 | 7.760 | 50.572 | 12.93 | 8.02 | n.a. |
| **Total:** | | | **60.024** | **630.914** | **100.00** | **100.00** | |

**Figure S3**. HPLC profiles of the protected MultiTASQ.

### MultiTASQ synthesis – Step 5:

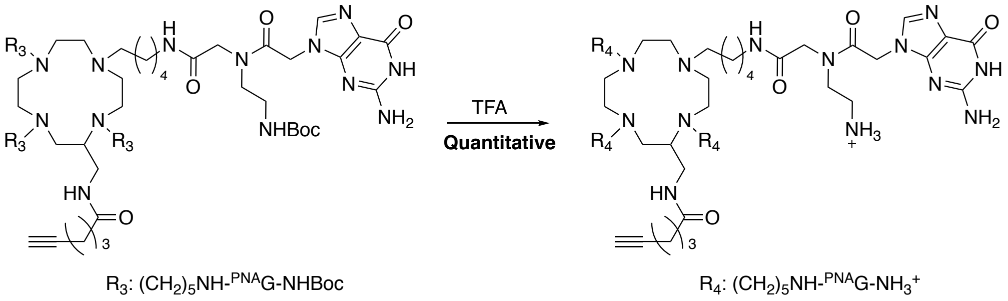


The protected MultiTASQ (1.95 mg, 0.9 nmol) was dissolved in TFA (100 μL) and the complete deprotection was assessed by HPLC-MS. The mixture was diluted in water and the compound was lyophilized to achieve MultiTASQ (2 mg, 0.9 nmol, yield 100%). ESI-HRMS: $[M+H]^{2+}$ m/z = 901.019 (calcd for $C_{79}H_{125}N_{37}O_{13}$: 901.020). HPLC-MS characterization (*Method A*): retention time = 0.32 min; purity: >97% at 201 nm, >93% at 280 nm; m/z = 901.7 $[M+H]^{2+}$ (cf. Figure S4). Due to its intrinsic very high polarity, the purity of MultiTASQ was further assessed by HPLC using an ACCUCORE column; retention time = 3.67 min; purity >96%; *Method C* (0% to 20% MeCN/$H_2O$, in 4 min and then from 20% to 100% MeCN/$H_2O$ + 0.1% formic acid (FA) in 2.6 min) (cf. Figures S4 and S5).

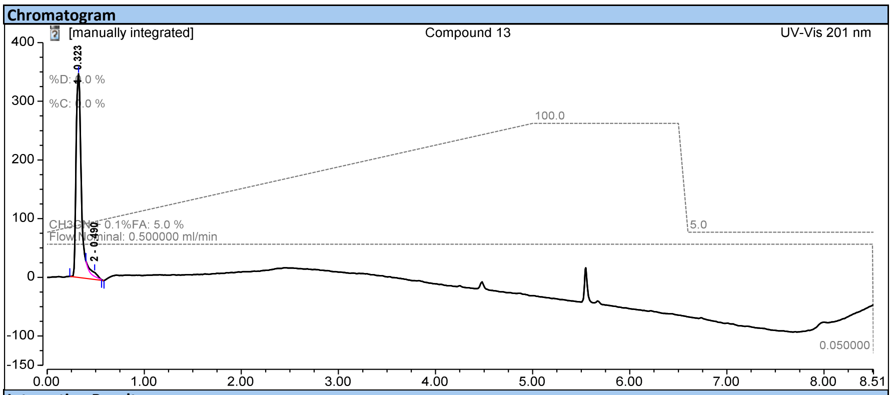


**Integration Results**

| No. | Peak Name | Retention Time min | Area mAU*min | Height mAU | Relative Area % | Relative Height % | Amount n.a. |
|---|---|---|---|---|---|---|---|
| 1 | | 0.323 | 21.935 | 345.415 | 97.11 | 98.09 | n.a. |
| 2 | | 0.490 | 0.652 | 6.715 | 2.89 | 1.91 | n.a. |
| **Total:** | | | **22.587** | **352.130** | **100.00** | **100.00** | |

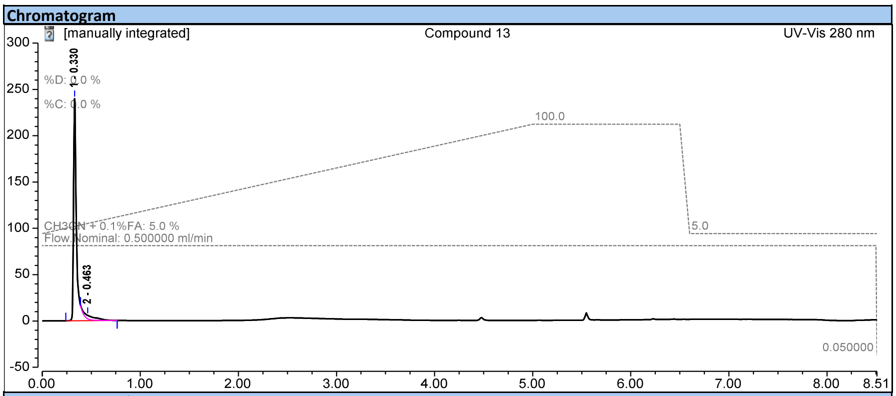


Integration Results

| No. | Peak Name | Retention Time min | Area mAU*min | Height mAU | Relative Area % | Relative Height % | Amount n.a. |
|---|---|---|---|---|---|---|---|
| 1 | | 0.330 | 8.430 | 239.683 | 93.22 | 98.50 | n.a. |
| 2 | | 0.463 | 0.613 | 3.652 | 6.78 | 1.50 | n.a. |
| Total: | | | 9.044 | 243.335 | 100.00 | 100.00 | |

**Figure S4**. HPLC and HPLC-MS profiles of MultiTASQ

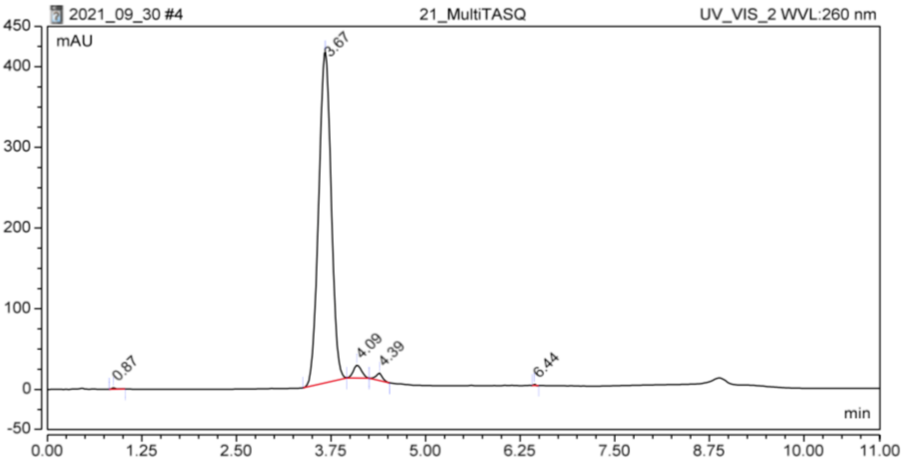


**Figure S5**. HPLC profile (Accucore) of MultiTASQ

### azMultiTASQ synthesis – Step 1:

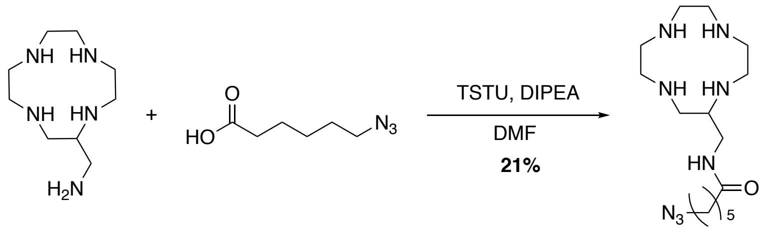


To a solution of 6-azido-hexanoic acid (414 mg, 2.64 mmol, 0.9 equiv.) and DIPEA (533 µL, 3.21 mmol, 1.1 equiv.) in DMF (2 mL) was added TSTU (995.7 mg, 3.21 mmol, 1.1 equiv.) and the solution was allowed to stir 1 h until complete reaction of the starting material (assessed by HPLC-MS). To a solution of AMC (588.9 mg, 2.92 mmol, 1 equiv.) in DMF (250 mL) at room temperature was added dropwise the latter reaction mixture containing N-hydroxylsuccimide-6-azido-hexanoate (1 mL/3 hrs). The reaction was carefully monitored by HPLC-MS and the addition was stopped as soon as the *bis*-substituted compound was observed. The solution was then concentrated under vacuum, water was added and the mixture purified by semi-preparative RP-HPLC in a $H_2O$/MeCN + 0.1% TFA mixture (gradient of 5 to 50% over 35 minutes, r.t: 13 minutes). After evaporation of the solvents, the acylated AMC was obtained (416.0 mg, 0.61 mmol, 21% chemical yield). $^{1}$H NMR: (500 MHz, $D_2O$): δ 3.31 (d, *J* = 5.4 Hz, 2H), 3.21 (t, *J* = 6.8 Hz, 2H), 3.17 – 2.81 (m, 15H), 2.19 (t, *J* = 7.4 Hz, 2H), 1.51 (m, 4H), 1.27 (m, 4.1 Hz, 2H). $^{13}$C NMR: (126 MHz, $D_2O$): δ 177.8, 119.7, 117.4, 115.1, 112.8, 51.0, 44.3, 43.9, 42.7, 42.1, 39.0, 35.4, 27.6, 25.5, 24.7. MALDI-TOF : $[M+H]^+$ m/z = 341.33 (calcd. for $C_{15}H_{33}N_8O$: 341.27). HPLC-MS characterization (Phenomenex Kinetex C18 column, 2.6µm, 2.1 × 50 mm) *via Method A* (from 5% to 100% MeCN/$H_2O$+0.1% formic acid (FA) in 7 min): retention time = 0.400min; purity: >92% at 201 nm, >93% at 214 nm; m/z = 341.6 $[M+H]^+$ (cf. Figure S6).

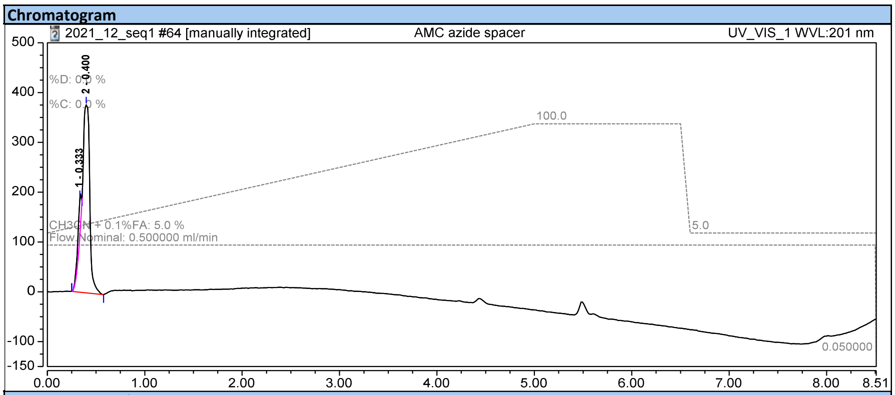


**Integration Results**

| No. | Peak Name | Retention Time min | Area mAU*min | Height mAU | Relative Area % | Relative Height % | Amount n.a. |
|---|---|---|---|---|---|---|---|
| 1 | | 0.333 | 2.993 | 77.108 | 7.74 | 16.98 | n.a. |
| 2 | | 0.400 | 35.666 | 377.123 | 92.26 | 83.02 | n.a. |
| **Total:** | | | **38.659** | **454.231** | **100.00** | **100.00** | |

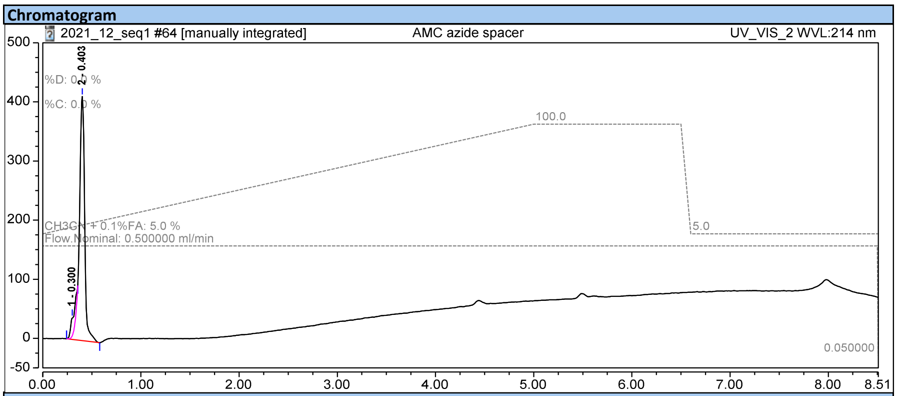


**Integration Results**

| No. | Peak Name | Retention Time min | Area mAU*min | Height mAU | Relative Area % | Relative Height % | Amount n.a. |
|---|---|---|---|---|---|---|---|
| 1 | | 0.300 | 1.942 | 30.994 | 6.95 | 6.98 | n.a. |
| 2 | | 0.403 | 25.991 | 413.087 | 93.05 | 93.02 | n.a. |
| Total: | | | 27.932 | 444.081 | 100.00 | 100.00 | |

**Figure S6**. HPLC profiles of the azido-AMC

### azMultiTASQ synthesis – Step 2:

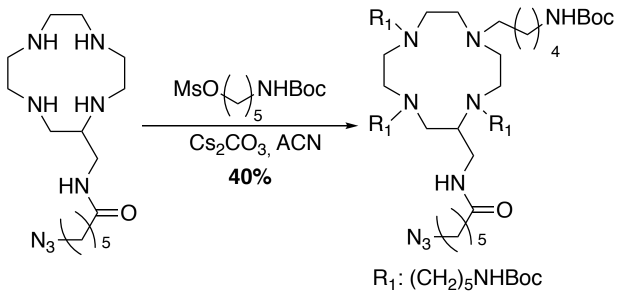


To a solution of the previously prepared acylated AMC (145.7 mg, 0.21 mmol, 1 equiv.) in MeCN (2.5 mL) was added 5-((*tert*-butoxycarbonyl)amino)pentyl methanesulfonate (576.6 mg, 2.1 mmol, 10 equiv.) and $Cs_2CO_3$ (668.3 mg, 2.1 mmol, 10 equiv.). The solution was heated at 50 °C and stirred for 72 hours, being monitored by HPLC-MS. After complete conversion, the crude was filtered and concentrated under vacuum. The residue was then purified by semi-preparative RP-HPLC with a $H_2O$/MeCN + 0.1% TFA mixture as mobile phase (gradient of 5 to 60 % over 50 minutes, r.t: 22 minutes). The Boc-protected compound was isolated as a white solid (92.4 mg, 0.085, mmol, 40 % chemical yield). $^1$H NMR: (500 MHz, Methanol-$d_4$) δ 3.73 – 3.25 (m, 6H), 3.22 (s, 4H), 3.19 – 3.02 (m, 9H), 2.96 (m, 10H), 2.75 (t, $J$ = 36.3 Hz, 7H), 2.15 (t, $J$ = 7.5 Hz, 2H), 1.84 – 1.49 (m, 10H), 1.48 – 1.38 (m, 9H), 1.34 (s, 42H), 1.26 – 1.17 (m, 4H). MALDI-TOF : $[M+H]^+$ m/z = 1081.99 (calcd. for $C_{55}H_{109}N_{12}O_9$: 1081.84). HPLC-MS characterization (*Method A)*: retention time = 4.270min; purity: >94% at 201 nm, >89% at 214 nm; m/z = 1082.6 $[M+H]^+$ (cf. Figure S7).

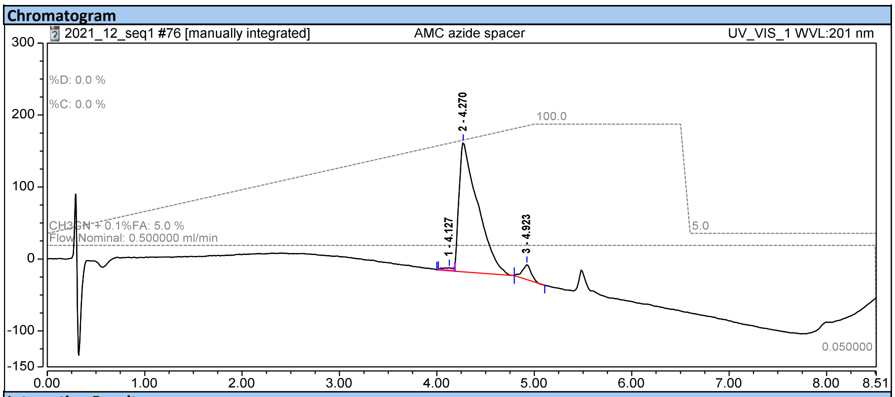


**Integration Results**

| No. | Peak Name | Retention Time min | Area mAU*min | Height mAU | Relative Area % | Relative Height % | Amount n.a. |
|---|---|---|---|---|---|---|---|
| 1 | | 4.127 | 0.384 | 3.246 | 0.90 | 1.59 | n.a. |
| 2 | | 4.270 | 40.303 | 179.567 | 94.10 | 88.22 | n.a. |
| 3 | | 4.923 | 2.145 | 20.724 | 5.01 | 10.18 | n.a. |
| **Total:** | | | **42.832** | **203.537** | **100.00** | **100.00** | |

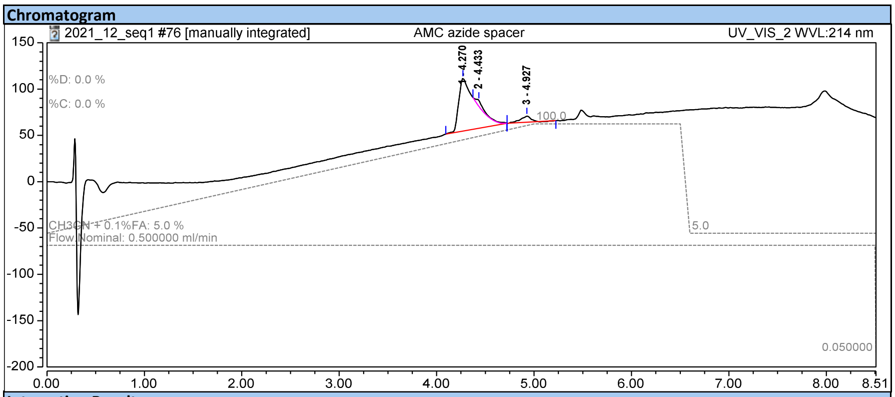


**Integration Results**

| No. | Peak Name | Retention Time min | Area mAU*min | Height mAU | Relative Area % | Relative Height % | Amount n.a. |
|---|---|---|---|---|---|---|---|
| 1 | | 4.270 | 11.520 | 56.849 | 89.24 | 81.25 | n.a. |
| 2 | | 4.433 | 0.630 | 6.469 | 4.88 | 9.24 | n.a. |
| 3 | | 4.927 | 0.759 | 6.652 | 5.88 | 9.51 | n.a. |
| **Total:** | | | **12.908** | **69.970** | **100.00** | **100.00** | |

**Figure S7**. HPLC profiles of the Boc-protected compound.

### $^{az}$MultiTASQ synthesis – Steps 3 and 4:

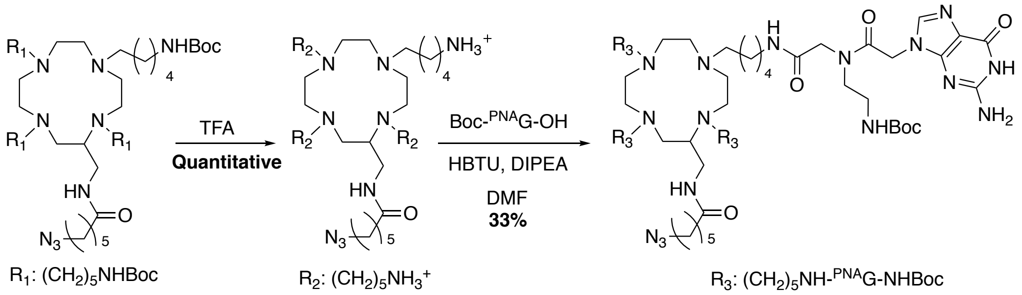


The previously prepared Boc-protected compound (50.6 mg, 0.042 mmol, 1 equiv.) was stirred with 500 μL of TFA for 1 h to deprotect the amines. After evaporation of the TFA (completion of the deprotection assessed by HPLC-MS), the deprotected compound was used without purification. In another round bottom flask, Boc-$^{PNA}$G (76.1 mg, 0.186 mmol, 4.4 equiv.), HBTU

(70.5 mg, 0.19 mmol, 4.4 equiv.) were dissolved in DMF (1 mL), and DIPEA was added (43 µL, 6 equiv.) The mixture was added to the solution containing previously prepared compound (50.6 mg, 0.042 mmol, 1 equiv.) and DIPEA (3 µL, 1 equiv.) in DMF (1 mL). The mixture was stirred at RT for 3 h  and the completion of the four couplings assessed by HPLC-MS. The solution was then concentrated under vacuum, diluted in a mixture of water and MeCN (50/50, 2mL), and purified by semi-preparative RP-HPLC in a $H_2O$/MeCN + 0.1% TFA mixture (gradient of 5 to 15% over 5 minutes then from 15 to 65% over 50 minutes, column *A*, r.t: 37 minutes). After evaporation of the solvents, the protected $^{az}$MultiTASQ was obtained (33.6 mg, 0.015 mmol, 33% chemical yield). ESI-HRMS**:** $[M+H]^{2+}$ m/z = 1123.641 (calcd. for $C_{99}H_{161}N_{40}O_{21}$: 1123.641). HPLC-MS characterization (*Method A)*: retention time = 0.400min; purity: >98% at 201 nm, >98% at 214 nm; >97% at 280 nm m/z = 1124.8 $[M+H]^{2+}$ (cf. Figure S8).

**Chromatogram**

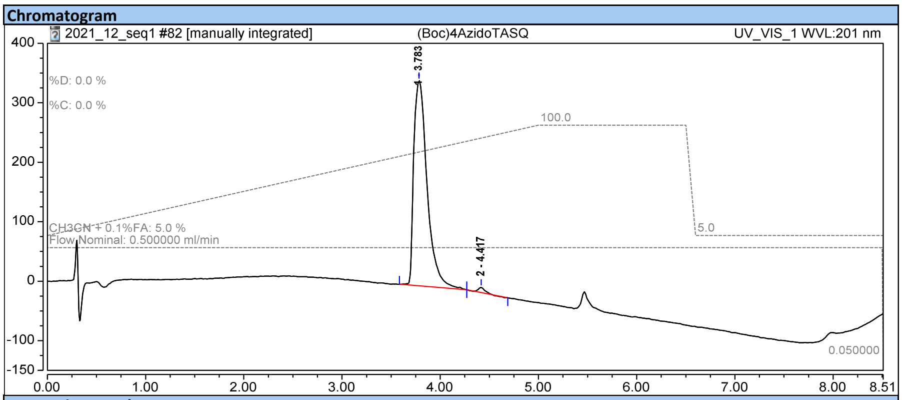


**Integration Results**

| No. | Peak Name | Retention Time min | Area mAU*min | Height mAU | Relative Area % | Relative Height % | Amount n.a. |
|---|---|---|---|---|---|---|---|
| 1 | | 3.783 | 54.214 | 345.001 | 98.86 | 97.55 | n.a. |
| 2 | | 4.417 | 0.627 | 8.655 | 1.14 | 2.45 | n.a. |
| **Total:** | | | **54.842** | **353.656** | **100.00** | **100.00** | |

**Chromatogram**

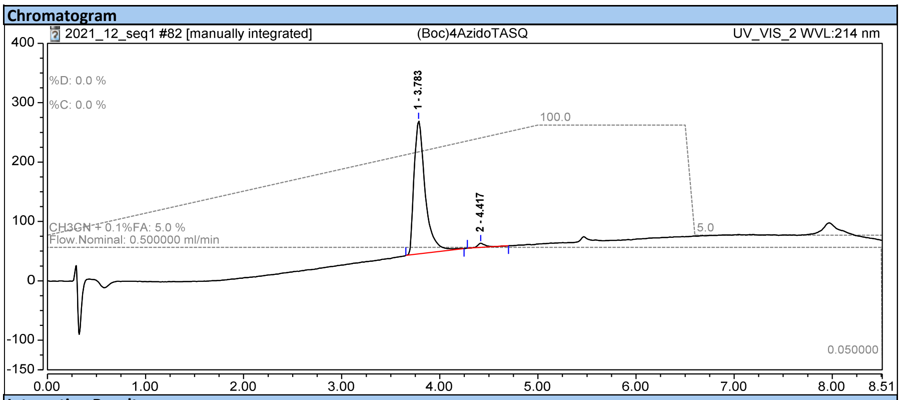


**Integration Results**

| No. | Peak Name | Retention Time min | Area mAU*min | Height mAU | Relative Area % | Relative Height % | Amount n.a. |
|---|---|---|---|---|---|---|---|
| 1 | | 3.783 | 29.495 | 223.833 | 98.10 | 97.01 | n.a. |
| 2 | | 4.417 | 0.573 | 6.909 | 1.90 | 2.99 | n.a. |
| **Total:** | | | **30.068** | **230.743** | **100.00** | **100.00** | |

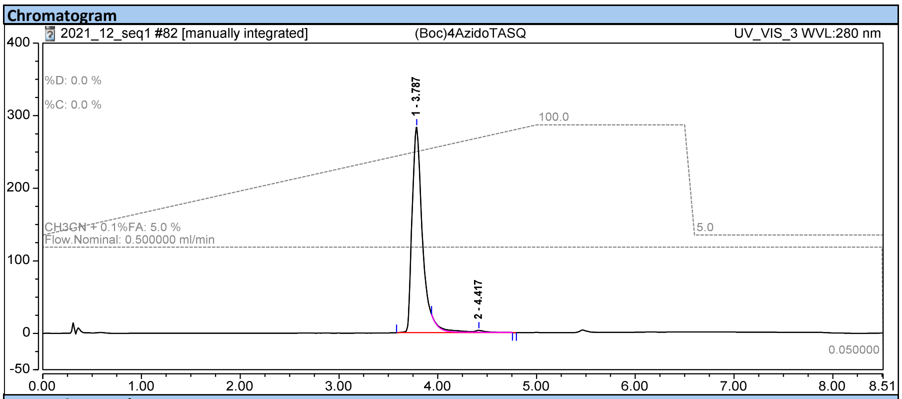


**Integration Results**

| No. | Peak Name | Retention Time min | Area mAU*min | Height mAU | Relative Area % | Relative Height % | Amount n.a. |
|---|---|---|---|---|---|---|---|
| 1 | | 3.787 | 34.594 | 282.768 | 97.59 | 99.08 | n.a. |
| 2 | | 4.417 | 0.856 | 2.640 | 2.41 | 0.92 | n.a. |
| **Total:** | | | **35.450** | **285.408** | **100.00** | **100.00** | |

**Figure S8**. HPLC profiles of the protected $^{az}$MultiTASQ.

### $^{az}$MultiTASQ synthesis – Step 5:

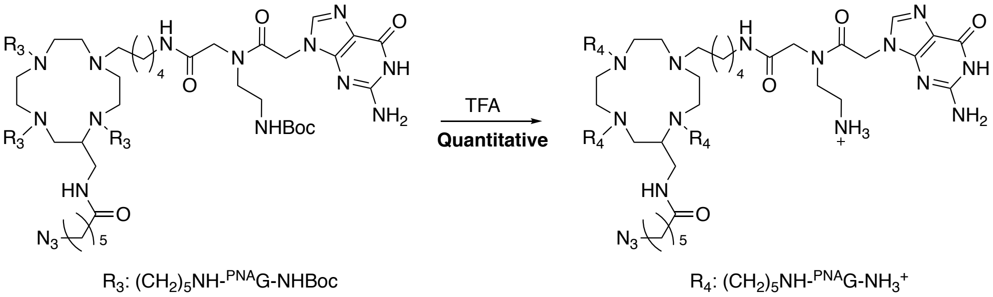


The protected $^{az}$MultiTASQ (1.95 mg, 0.9 µmol) was dissolved in TFA (100 µL) and the complete deprotection was assessed by HPLC. The mixture was diluted in water and the compound was lyophilized to afford $^{az}$MultiTASQ. (2 mg, 0.9 µmol, chemical yield 100%). ESI-HRMS: $[M+H]^{3+}$ m/z = 616.026 (calcd. for $C_{79}H_{129}N_{40}O_{13}$: 616.026). HPLC-MS characterization (*Method A*): retention time = 0.38 min; purity: >97% at 214 nm, >98% at 280 nm; m/z = 924.3 $[M+H]^{2+}$ (cf. Figure S9). Due to its intrinsic very high polarity, the purity of MultiTASQ was further assessed by HPLC using an ACCUCORE column, retention time = 3.84 min; purity >90%; *Method C* (0% to 20% MeCN/$H_2O$, in 4 min and then from 20% to 100% MeCN/$H_2O$ + 0.1% formic acid (FA) in 2.6 min) (cf. Figures S9 and S10).

**Chromatogram**

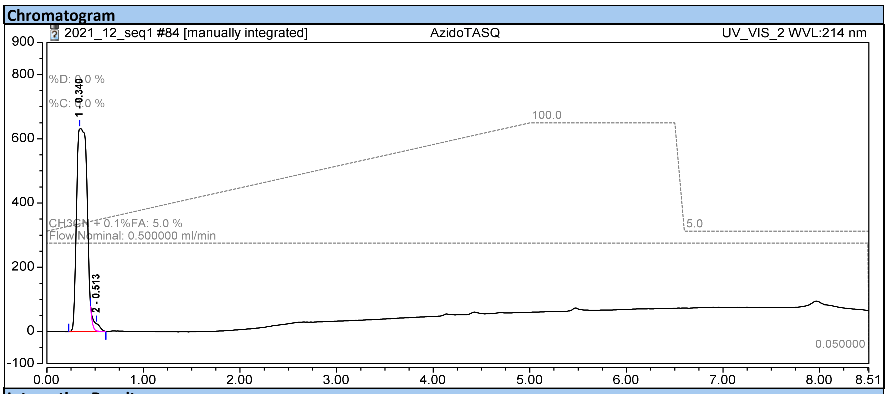


**Integration Results**

| No. | Peak Name | Retention Time<br>min | Area<br>mAU*min | Height<br>mAU | Relative Area<br>% | Relative Height<br>% | Amount<br>n.a. |
|---|---|---|---|---|---|---|---|
| 1 | | 0.340 | 80.331 | 633.718 | 97.58 | 96.64 | n.a. |
| 2 | | 0.513 | 1.989 | 22.031 | 2.42 | 3.36 | n.a. |
| **Total:** | | | **82.320** | **655.749** | **100.00** | **100.00** | |

**Chromatogram**

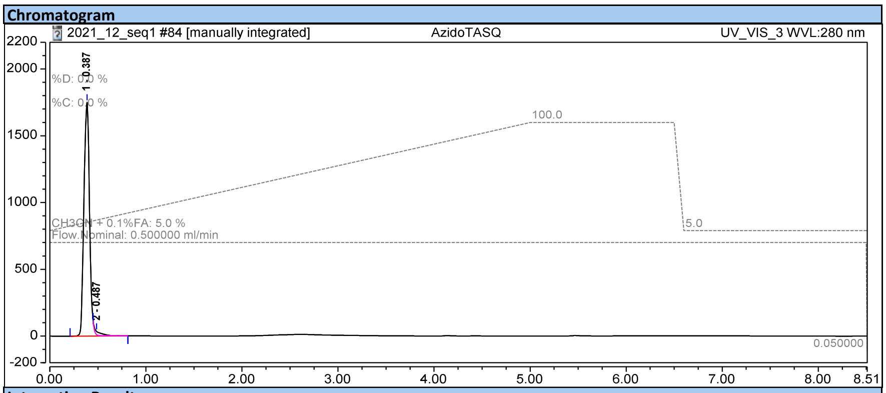


**Integration Results**

| No. | Peak Name | Retention Time<br>min | Area<br>mAU*min | Height<br>mAU | Relative Area<br>% | Relative Height<br>% | Amount<br>n.a. |
|---|---|---|---|---|---|---|---|
| 1 | | 0.387 | 119.673 | 1751.006 | 98.04 | 98.67 | n.a. |
| 2 | | 0.487 | 2.389 | 23.689 | 1.96 | 1.33 | n.a. |
| **Total:** | | | **122.062** | **1774.695** | **100.00** | **100.00** | |

**Figure S9**. HPLC profiles of $^{az}$MultiTASQ.

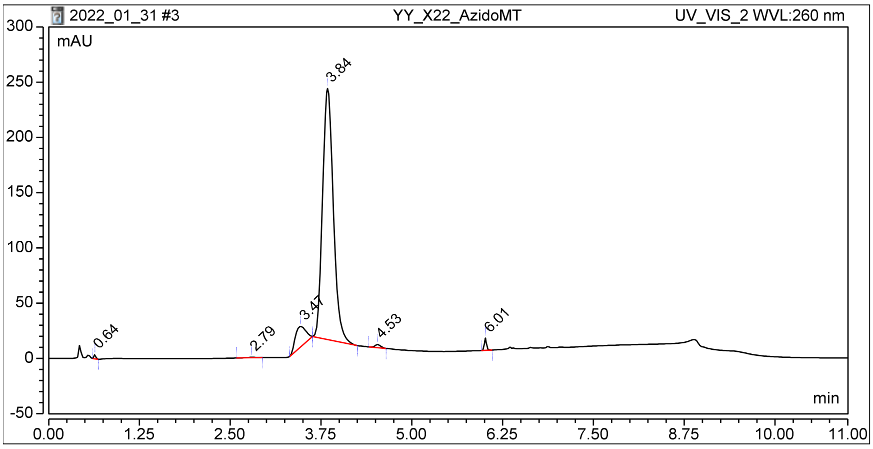


**Figure S10**. HPLC profile (Accucore) of $^{az}$MultiTASQ.

**II. Oligonucleotides.**

The lyophilized DNA/RNA strands (purchased from Eurogentec) were firstly diluted in deionized water (18.2 MΩ.cm resistivity) at 500 µM. All DNA structures were prepared in a Caco.K buffer, comprised of 10 mM lithium cacodylate buffer (pH 7.2) plus 10 mM KCl/90 mM LiCl. Structures were prepared by mixing 40 µL of the constitutive strand (500 µM) with 8 µL of a lithium cacodylate buffer solution (100 mM, pH 7.2), plus 8 µL of a KCl/LiCl solution (100 mM/900 mM) and 24 µL of water. The final concentrations, theoretically 250 µM, was determined through a dilution to 1 µM theoretical concentration (expressed in motif concentration; *i.e.*, 4 µL in 996 µL water), through UV spectral analysis at 260 nm (after 5 min at 90 °C) with the molar extinction coefficient values provided by the manufacturer. The higher-order DNA/RNA structures were folded by heating the previously prepared solutions at 90 °C for 5 min, cooling them directly on ice (several hours) and then storing them at least overnight at 4 °C.

**Table S1. Doubly labeled oligonucleotides** (FRET-melting assay)

| **Nature** | **Name** | **Sequence** |
|---|---|---|
| **DNA** | F21T | FAM-d[$^{5'}$GGGTTAGGGTTAGGGTTAGGG$^{3'}$]-TAMRA |
| **DNA** | F-myc-T | FAM-d[$^{5'}$GAGGGTGGGGAGGGTGGGGAAG$^{3'}$]-TAMRA |
| **RNA** | F-TERRA-T | FAM-r[$^{5'}$GGGUUAGGGUUAGGGUUAGGG$^{3'}$]-TAMRA |
| **RNA** | F-VEGF-T | FAM-r[$^{5'}$GGAGGAGGGGAGGAGGA$^{3'}$]-TAMRA |
| **RNA** | F-NRAS-T | FAM-r[$^{5'}$GGGAGGGGCGGGUCUGGG$^{3'}$]-TAMRA |

**Table S2. Labeled oligonucleotides** ($K_D$ measurements and fluorescence pull-down exp.)

| **Nature** | **Name** | **Sequence** |
|---|---|---|
| **DNA** | Cy5-Myc | Cy5-d[$^{5'}$GAGGGTGGGGAGGGTGGGGAAG$^{3'}$] |
| **DNA** | F-22AG | FAM-d[$^{5'}$AGGGTTAGGGTTAGGGTTAGGG$^{3'}$] |
| **DNA** | F-Myc | FAM-d[$^{5'}$GAGGGTGGGGAGGGTGGGGAAG$^{3'}$] |
| **DNA** | F-duplex | FAM-d[$^{5'}$TATAGCTATATTTTTTTATAGCTATA$^{3'}$] |
| **RNA** | F-TERRA | FAM-r[$^{5'}$GGGUUAGGGUUAGGGUUAGGG$^{3'}$] |
| **RNA** | F-VEGF | FAM-r[$^{5'}$GGAGGAGGGGAGGAGGA$^{3'}$] |
| **RNA** | F-NRAS | FAM-r[$^{5'}$ GGGAGGGGCGGGUCUGGG $^{3'}$] |

**Table S3. Unlabeled oligonucleotides** (qPCR pull-down exp. and G4RP protocol)

| **Nature** | **Name** | **Sequence** |
|---|---|---|
| **DNA** | G4-strand | d[5'TAGCCATTCAGCCGTAACAGGCAGTGGAAGAGAGACAGACAGGGCAGGGCAGGGCAGGGCAGTACAGTAGAACCTAATGGTGTTTGATGGTATCTAA3'] |
| **DNA** | NRAS forward | d[5'ATGACTGAGTACAAACTGGTGGT 3'] |
| **DNA** | NRAS reverse | d[5'CATGTATTGGTCTCTCATGGCAC 3'] |
| **DNA** | VEGF forward | d[5'CCTTGCCTTGCTGCTCTACC 3'] |
| **DNA** | VEGF reverse | d[5'AGATGTCCACCAGGGTCTCG 3'] |

**Table S4. Used buffers**

| **Nature** | **Composition** |
|---|---|
| CacoK1 | 10 mM lithium cacodylate buffer (pH 7.2) + 1 mM KCl/99 mM LiCl |
| CacoK10 | 10 mM lithium cacodylate buffer (pH 7.2) + 10 mM KCl/90 mM LiCl |
| TrisHCl/Triton | TrisHCl (pH 7.2) + 150mM KCl and 0.05 v/v % Triton X-100 |
| TrisHCl/$MgCl_2$ | 20 mM TrisHCl buffer (pH 7.2), 1 mM KCl, 99 mM LiCl and 10 mM $MgCl_2$ |
| DEPC-$H_2O$ | 1 mL DEPC + 1000 mL of dd$H_2O$ |
| DEPC-PBS | Dilute 10X PBS stock in DEPC-$H_2O$ |
| Fixing buffer | 250 mM HEPES KOH pH 7.5, 500 mM NaCl, 5 mM EDTA pH 8, 2.5 mM EGTA pH 8 in DEPC-$H_2O$ |
| G4RP Buffer | 150 mM KCl, 25 mM Tris pH 7.4, 5 mM EDTA, 0.5 mM DTT, 0.5% NP40 (tergitol), pH 7.4, 0.1 U RNAse OUT in DEPC-$H_2O$ |

### III. Additional *in vitro* results

We hereby control whether 1) the biotinylation of MultiTASQs affects their G4-stabilization properties by FRET-melting; 2) MultiTASQs do interact efficiently with N-RAS G4, first by FRET-melting and then by fluorescence pull-down, and 3) the presence of the copper was detrimental for the use of the clicked MultiTASQ: to this end, a solution of the biotinylated and metalated TASQ was split in two, half of this solution was demetallated ($Na_2S$ treatment), the other half was not, and the resulting TASQs for by fluorescence pull-down experiments.

**A.** FRET-melting controls

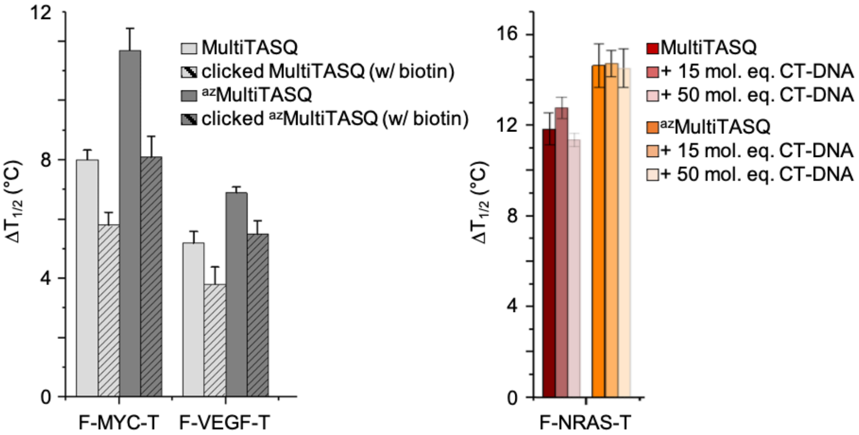


| | F-MYC-T | F-VEGF-T |
|---|---|---|
| MultiTASQ | 8.0 ± 0.3 | 5.2 ± 0.4 |
| Clicked MultiTASQ | 5.8 ± 0.4 | 3.8 ± 0.6 |
| $^{az}$MultiTASQ | 11.7 ± 0.7 | 6.9 ± 0.2 |
| Clicked $^{az}$MultiTASQ | 8.1 ± 0.7 | 5.5 ± 0.5 |

| | F-NRAS-T |
|---|---|
| MultiTASQ | 11.8 ± 0.7 |
| +50 mol. equiv. CT-DNA | 11.3 ± 0.3 |
| $^{az}$MultiTASQ | 14.6 ± 1.1 |
| +50 mol. equiv. CT-DNA | 14.5 ± 0.8 |

**B.** Fluorescence pull-down controls

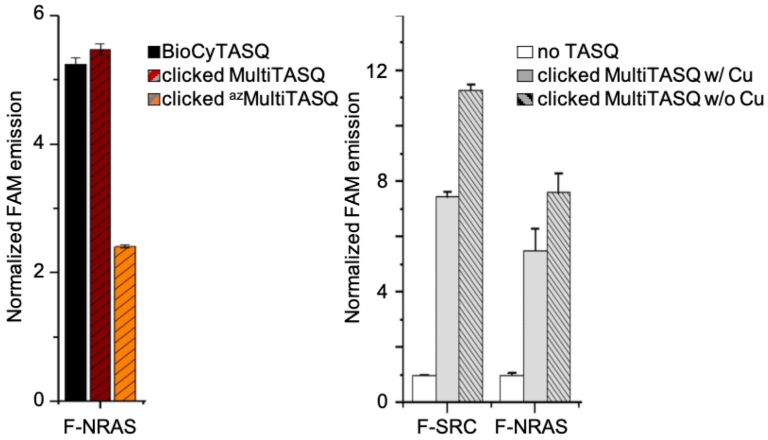


| | F-NRAS |
|---|---|
| BioCyTASQ | 5.2 ± 0.9 |
| Clicked MultiTASQ | 5.5 ± 0.8 |
| Clicked $^{az}$MultiTASQ | 2.4 ± 0.3 |

| | F-SRC | F-NRAS |
|---|---|---|
| MultiTASQ w/ copper | 7.43 ± 0.2 | 5.5 ± 0.8 |
| MultiTASQ w/o copper | 11.3 ± 0.2 | 7.6 ± 0.7 |

**Figure S11**. FERT-melting and fluorescence pull-down control experiments.
